# Brain-specific *Gata4* downregulation in *Greywick* female mice models the metabolic subtype of polycystic ovary syndrome

**DOI:** 10.1101/2024.05.13.593880

**Authors:** Sherin A. Nawaito, Mostafa Esmael, Ouliana Souchkova, Tatiana Cardinal, Guillaume Bernas, Karl-F. Bergeron, Fanny Gayda, Francis Bergeron, Marie-France Bouchard, Xiang Zhou, Luisina Ongaro, Daniel J. Bernard, Jacob Short, Susan Wray, Robert S. Viger, Catherine Mounier, Nicolas Pilon

**Affiliations:** Molecular Genetics of Development Laboratory, Department of Biological Sciences, Université du Québec à Montréal, Montreal H3C 3P8, Quebec, Canada; Centre d’Excellence en Recherche sur les Maladies Orphelines – Fondation Courtois, Université du Québec à Montréal, Montreal H2X 3Y7, Canada; Department of Physiology, Suez Canal University, Ismailia, Egypt; Lipid Metabolism Laboratory, Department of Biological Sciences, Université du Québec à Montréal, Montreal H3C 3P8, Canada; Reproduction, Mother and Child Health, Centre de recherche du CHU de Québec-Université Laval, Québec GIV 4G2, Canada; Department of Pharmacology and Therapeutics, McGill University, Montreal H3G 1Y6, Canada; Cellular and Developmental Neurobiology Section, National Institute of Neurological Disorders and Stroke, Bethesda, MD 20814, United States of America; Department of Obstetrics, Gynecology, and Reproduction, Université Laval, and Centre de recherche en reproduction, développement et santé intergénérationnelle, G1K 7P4, Canada; Department of Pediatrics, Université de Montréal, Montreal H3T 1C5, Canada

**Keywords:** Polycystic ovary syndrome, GATA4, mouse model, fertility, metabolism, hypothalamus

## Abstract

Polycystic ovary syndrome (PCOS) is a heterogenous disorder characterized by reproductive and metabolic abnormalities. PCOS etiology remains poorly understood, although the hypothalamus is suspected to play a central role in many cases. Human genetic studies have also shown an association with the transcription factor-coding gene *GATA4*, but without providing a functional link. Here, we show that adult *Greywick* female mice may bridge this gap. These mice phenocopy PCOS with partial penetrance, due to serendipitous insertion of a *Gata4* promoter-driven transgene in a strong enhancer region. Resulting robust transgene expression in subsets of hypothalamic neurons and glia impairs endogenous *Gata4* expression, resulting in misexpression of genes linked to the control of fertility and food intake. We also show that this previously overlooked role of GATA4 in the hypothalamus can be replicated by conditional knockout approaches. Overall, this study sheds light not only on PCOS etiology but also on the role played by GATA4 in the central control of reproduction.

## INTRODUCTION

Polycystic ovary syndrome (PCOS) is a multifactorial disorder and the most common diagnosis of female infertility worldwide ^1, 2, 3^. Clinical presentation is heterogeneous, with the presence of ovarian follicular cysts not even necessary for reaching diagnosis ^1, 4^. According to the consensus-based 2003 Rotterdam criteria and 2023 international evidence-based guidelines, only two out the following three main features are needed to establish diagnosis: 1) oligo/anovulation, 2) clinical/biochemical hyperandrogenism, and 3) either multiple ovarian follicular cysts or elevated levels of anti-Müllerian hormone (AMH) ^1, 2, 3, 5^. In addition, PCOS is often associated with mood disorders as well as metabolic anomalies, including obesity, dyslipidemia, and glucose/insulin resistance, which are all risk factors of cardiovascular complications ^1, 3, 6^. Accordingly, PCOS is currently classified as an endocrine spectrum disorder, with a “reproductive” subtype at one end and a “metabolic” subtype at the other ^4, 7, 8, 9, 10^. The reproductive subtype is notably characterized by higher levels of luteinizing hormone (LH) and relatively normal body mass index and insulin levels, whereas the metabolic subtype is still characterized by elevated LH levels (albeit not as high) combined with high body mass index and insulin levels ^7^.

Regardless of the clinical subtype, androgen excess appears to play a critical role in PCOS pathogenesis ^1, 3, 5^. The major role of androgens is made evident in several PCOS animal models that are induced with either the non-aromatizable androgen dihydrotestosterone (DHT), the “weak” adrenal androgen dehydroepiandrosterone (DHEA), or testosterone propionate (TP) ^11, 12, 13, 14, 15^. Yet, in a real-life PCOS context, increased androgen levels appear to be the consequence of hypothalamic resistance to negative feedback signals from ovary-derived estrogen and progesterone ^5, 16, 17, 18^. This hypothalamic resistance leads to increased pulsatility of GnRH and LH secretion ^5, 19^, resulting in persistently high levels of LH that trigger increased testosterone production by follicular theca cells in the ovary ^5, 20^. High levels of circulating testosterone can then activate androgen receptor (AR) signaling in many PCOS-relevant tissues, with neuroendocrine circuits of the hypothalamus being a likely major site of action ^21^. In agreement with this mechanistic model, analysis of tissue-specific AR knockout mice revealed that most of the postnatal DHT effects as an inducer of PCOS-like traits are mediated by AR signaling in neurons rather than in ovaries ^22^.

In the metabolic PCOS subtype, hyperinsulinemia (secondary to insulin resistance in a highly metabolically active tissue like the liver, adipose tissue, or skeletal muscle) further enhances the vicious cycle of hypothalamus-pituitary-ovary axis dysfunction ^23, 24^. In the hypothalamus, overactive insulin signaling is believed to contribute to the increased release of GnRH/LH, either directly by acting on GnRH neurons (increasing *Gnrh1* transcription) ^25, 26^ and/or indirectly via the GnRH pulse generator KNDy (kisspeptin/neurokinin B/dynorphin) neurons ^27^. In addition, insulin excess acts directly on ovarian theca cells, thereby contributing to the increased secretion of androgens ^24, 28^. In turn, androgen excess can enhance insulin resistance by increasing visceral adiposity and levels of fatty acids in peripheral tissues, thus creating the other vicious cycle of insulin resistance-hyperinsulinemia-hyperandrogenism ^1, 23, 24^. Of note, accumulation of fatty acids in the ovary can also negatively impact follicle maturation and oocyte quality by increasing endoplasmic reticulum (ER) stress ^29, 30^ and mitochondrial dysfunction^31^.

PCOS has strong genetic roots, most likely with additional involvement of environmental factors like diet and lifestyle ^1, 3, 4, 32^. Based on several genome-wide association studies (GWAS), including a meta-analysis of >10,000 cases and >100,000 controls, >20 genetic loci have thus far been linked to PCOS ^7, 33, 34, 35, 36, 37, 38^. Yet, for most of the identified single nucleotide polymorphisms (SNPs), the mediating genes and/or their functional effects are unclear ^39^. One PCOS-associated SNP that has been replicated in many cohorts (rs804279:A>T) is located ∼6 kb downstream of the transcription factor-coding gene *GATA4* ^34, 36^, and appears to be linked with higher risk of metabolic dysfunction ^8, 34^. Another SNP (rs3729853:T>C/G) located in the gene body of *GATA4* (near the end of intron 5) has been recently associated with both lean and overweight/obese subgroups of bodyweight-stratified cases of PCOS ^38^. Moreover, the GATA4 protein is present in growing follicles (in both granulosa and theca cells) and surface epithelium ^40^ of postnatal murine ovaries, and its ovary-specific loss leads to subfertility/sterility associated with depletion of the follicle pool and formation of inclusion cysts ^41, 42^. However, these cysts are of epithelial origin, not follicular like in PCOS, and no signs of PCOS-associated metabolic dysfunction were reported in these conditional *Gata4* knockout mice ^41, 42^. Combined with the other observation that *GATA4* expression levels were unaffected in ovarian samples from a cohort of 28 women with PCOS ^43^, all these findings strongly suggest that a yet to be identified extra-ovarian role of GATA4 is involved.

Here, we report that adult female mice from a *Gata4*-reporter transgenic line initially designed for studies in developmental biology, phenocopy, with partial penetrance, the main diagnostic features and commonly associated metabolic anomalies of human PCOS. Detailed characterization of this mouse line strongly suggests that these female-specific phenotypes are due to perturbation of a previously overlooked role of GATA4 in the hypothalamus, rather than in ovaries. Moreover, by also analyzing conditional knockout mice, we provide definitive genetic evidence that female fertility relies on brain GATA4.

## RESULTS

### Serendipitous generation of a mouse model for *GATA4*-associated PCOS

As part of a project focused on the analysis of cis-regulatory elements of *Gata4*, we previously generated five transgenic reporter lines that are now used as tools to endogenously label Sertoli ^44, 45, 46^ and neural crest ^47, 48, 49, 50^ cell lineages with either green (GFP) or red (RFP) fluorescence during prenatal development. For all five lines, reporter gene expression is driven by the 5 kb proximal promoter of rat *Gata4* exon E1a, and transgenic sequences also include a co-injected *tyrosinase* minigene that serves as a transgenesis marker in the FVB/N albino background (via pigmentation rescue) ^51^. Only one of these mouse lines could be maintained as homozygous, which we named *Greywick* (*Gw*) in reference to its coat color (Fig.1A). This line also drew our attention because of the female-specific phenotypes combining subfertility (Fig.1B-C) and overweight (Fig.1D) in about a quarter of adults, reminiscent of the metabolic subtype of PCOS. As described in more detail in the next section, both phenotypes were found to be partially penetrant, and not to be systematically associated (Fig.1E). Subfertility and/or overweight were never observed at the heterozygous state – neither for this line nor any of the four other *Gata4*-reporter lines, which could only be maintained as heterozygotes because of X chromosome insertion or homozygosity-associated embryonic lethality ^44^. Moreover, these phenotypes were never observed in males. *Gw* male mice are as fertile (Fig.S1A) and as lean (Fig.S1B) as control FVB/N male mice of the same age.

**Figure 1.**
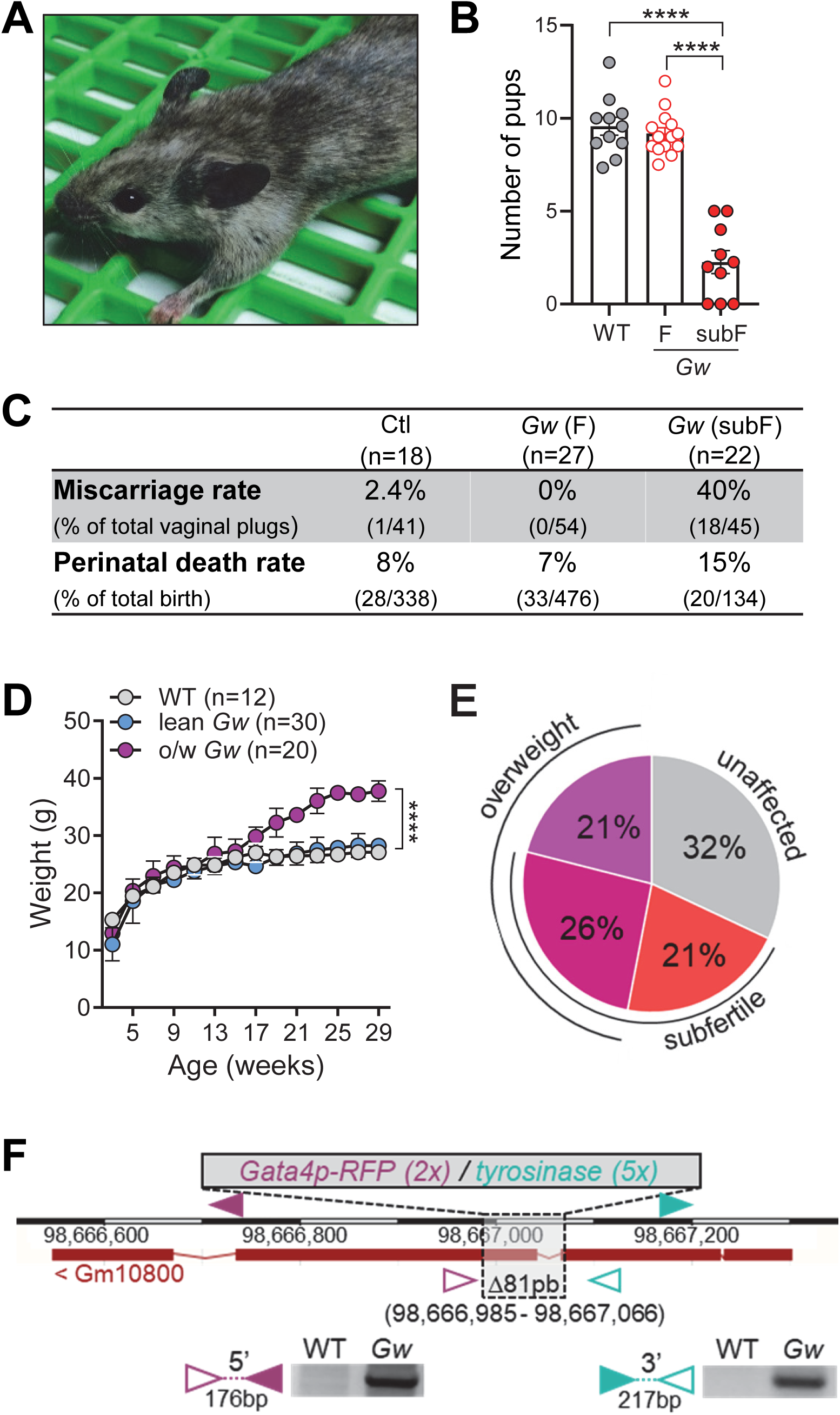
Subsets of *Greywick* female mice are subfertile and gain extraweight with age. **A**) Pigmentation pattern of so-named *Greywick* (*Gw*) mice in the FVB/N genetic background. **B)** Average number of pups per litter for adult *Gw* and WT females (2– to 6-month-old), after breeding with WT males (n=10-11 females per group, 2-3 litters per female). F, fertile; subF, subfertile. **C)** Comparative table of pregnancy complication-related parameters. **D)** Weight gain as a function of female age (*Gw vs* WT; n=12-30 females per group). **E)** Phenotypic distribution of subfertility and overweight in *Gw* female mice aged between 2 to 6 months (n=115 females). **F)** Overview of the *Gw* insertion site showing co-integration of *Gata4p[5kb]-RFP* (2 copies) and *tyrosinase* (5 copies) transgenes into an 81bp deletion (dashed box) within *Gm10800* on Chr.2 (adapted from the Ensembl website), and confirmatory PCR products for 5’ and 3’ genomic/transgenic boundaries using indicated color-coded primers. **** *p* < 0.0001; two-tailed Welch’s *t*-test (B) or two-way ANOVA with Dunnet’s post hoc test (C).

Using whole-genome sequencing and alignment on the GRCm38 reference genome (to which we also added our transgenic sequences), we localized the *Gw* transgenic insertion in *Gm10800* (predicted gene 10800) on Chr.2, band E1 (Fig.1F). Precise mapping proved difficult because of the presence of satellite-type repetitive sequences at this A/T-rich locus (64% of A/T content in *Gm10800*), requiring three independent rounds of whole-genome sequencing (including one using the Linked-Reads approach from 10X Genomics). Our mapping approach further allowed us to estimate the number of copies of each transgene (two for *Gata4p[5kb]-RFP vs.* five for *tyrosinase*), while anchored PCR combined to Sanger sequencing was used to definitively confirm the genomic/transgenic boundaries (Fig.1F).

To verify the possibility that *Gm10800* disruption could be responsible of the PCOS-like features in *Gw* mice, we generated an independent *Gm10800* knockout line in the same FVB/N background via CRISPR/Cas9 (Fig.2A). Adult females homozygous for this new *Gm10800*-null allele were indistinguishable from their WT littermates in terms of both fertility rate (Fig.2B) and body weight (Fig.2C). Of note, owing to the repetitive nature of this locus, our CRISPR/Cas9-based approach also yielded another mouse line with a larger deletion encompassing both *Gm10800* and its paralogous neighbour *Gm10801* (with 91% of DNA sequence identity). Adult females homozygous for this double knockout also had no overt problems of fertility (Fig.S1C) and/or body weight (Fig.S1D). This outcome was not surprising given what is now known about the genomic landscape of the *Gm10800/Gm10801* region, which also explains our prior inability to clone a full-length cDNA corresponding to the predicted open reading frame of *Gm10800*. Closer examination of this genomic landscape on both Ensembl and UCSC genome browsers clearly contradicts the initial GENCODE prediction in the GRCm38 reference genome that *Gm10800* is protein-coding – a prediction since retracted in the latest GRCm39 version. As shown on a screenshot of the UCSC genome browser (Fig.2D; zoomed out in Fig.S1E), predicted coding sequences of *Gm10800* in the GRCm38 reference genome contain hundreds of frameshifting indel variations (intolerable for any protein-coding gene), while multispecies sequence comparisons do not show any alignment (even with rat sequences). Instead, this region has all the hallmarks of a safe-harbor locus for robust transgene expression, with broad ATAC-seq peaks indicative of highly accessible chromatin (notably in brain tissues), ChIP-seq data for histone marks indicative of an active enhancer signature (H3K27ac+ H3K4me1+), and larger aggregate ChIP-seq data suggestive of a hotspot for transcription factor binding in a wide diversity of cell types (Fig.2D). Accordingly, cloned cDNAs that align with this region correspond to bidirectionally transcribed RNAs of variable lengths (Fig.S1E), and hence are most likely enhancer RNAs ^52^.

**Figure 2.**
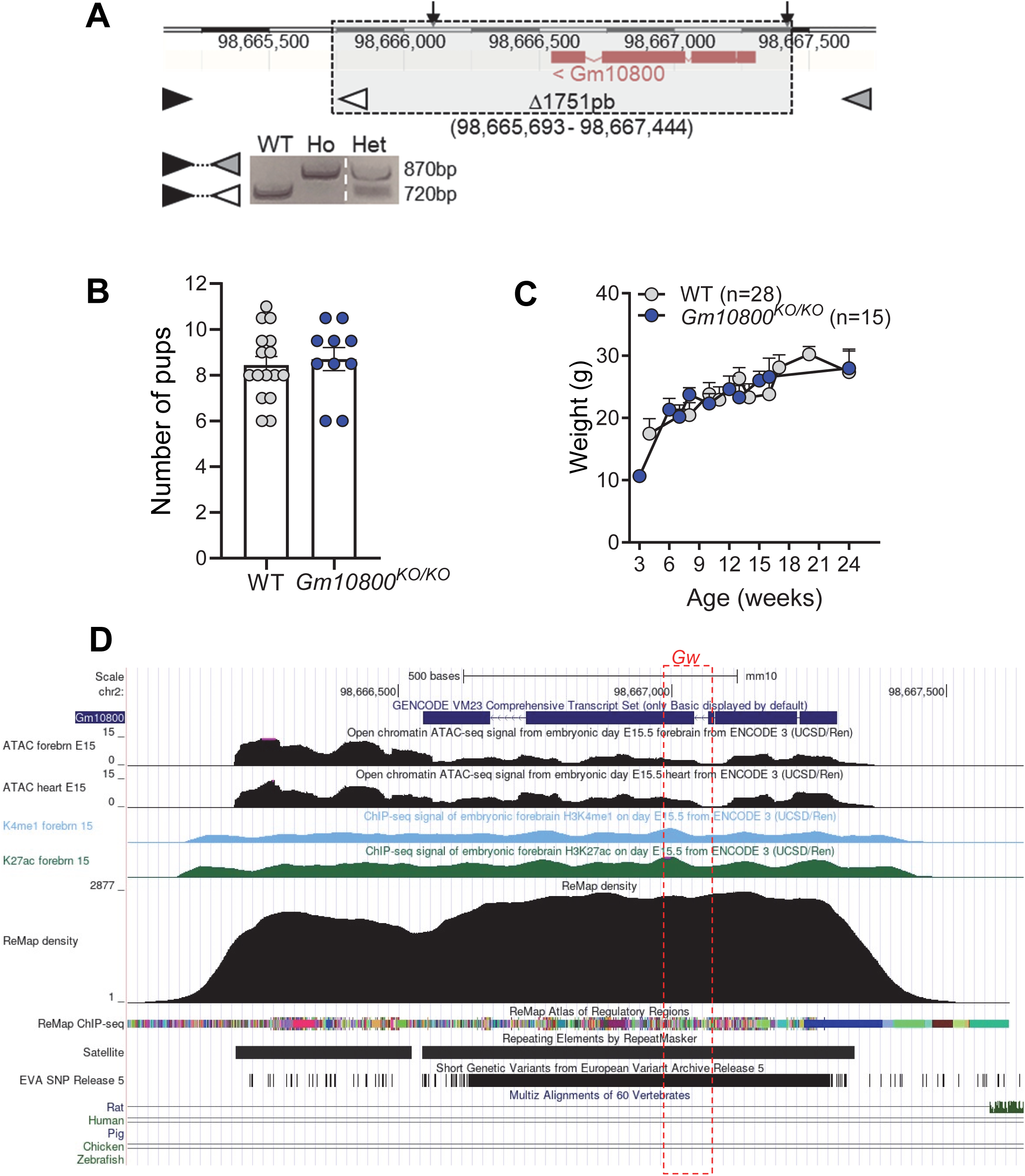
Disruption of *Gm10800* is not the cause of the *Gw* phenotypes. **A**) Overview of the *Gm10800* locus (adapted from the Ensembl website) showing position of sgRNAs (black arrows) and resulting 1751bp deletion after CRISPR/Cas9-mediated deletion and NHEJ repair (dashed box), and genotyping PCR products in mice using indicated color-coded primers. **B)** Average number of pups per litter for *Gm10800^KO/KO^* and WT females (2-to 6-month-old), after breeding with WT males (n=10-16 females per group, 2-3 litters per female). **C)** Weight gain as a function of female age (*Gm10800^KO/KO^ vs* WT; n=15-28 females per group). **D)** Screenshot from the UCSC genome browser showing key genomic features within and around *Gm10800*, with position of the *Gw* insertion site indicated by a dashed red box. ATAC-seq tracks show chromatin accessibility in forebrain and heart tissues at embryonic day (E)15, K4me1 and K27ac ChIP-seq tracks show the signature of an active enhancer in E15 forebrain, and ReMap tracks show aggregate ChIP-seq data for multiple transcription factors in a wide diversity of cell types. Other tracks at the bottom further show that this region is enriched in satellite repetitive sequences, highly tolerant to single nucleotide variations, and not conserved in any other species.

From all these mouse genetic data, we conclude that partially penetrant PCOS-like traits in adult homozygous *Gw* females (hereafter simply referred to as *Gw*, unless otherwise stated) are due to the combination of both the *Gata4* reporter transgene and its insertion site. The insertion of exogenous *Gata4* regulatory sequences within a particularly accessible and active enhancer region most likely perturbs endogenous *Gata4* expression via competition for a limited pool of transcriptional activators, as previously described in other contexts ^53, 54, 55^. Therefore, *Gw* females may represent a valuable model to better understand the association between *GATA4* and PCOS in humans ^34, 36, 38^.

### Overweight is not the cause of subfertility in *Gw* female mice

To validate our preliminary phenotypic observations, we systematically compared the fertility rate of adult *Gw* and wild-type (WT) control FVB/N females (aged between 2 to 6 months), both bred with fertile control males. Based on the average number of pups per litter for at least two pregnancies per female, *Gw* mice could be divided into fertile (≥7.5 pups per litter, like WT controls) and subfertile (≤5 pups per litter) subgroups representing 59% (13/22) and 41% (9/22) of animals in this cohort, respectively (Fig.1B). In another cohort of 4-month-old mice, we assessed the rates of miscarriage and perinatal death, finding that both parameters of pregnancy complication were much higher in sub-fertile *Gw* mice (Fig.1C), as they are in PCOS women ^3, 56^. Systematic comparison of the body weight of *Gw* and WT females, from birth until 29 weeks of age, also identified two subgroups of *Gw* mice, one designated as lean (with age-associated weight gain similar to WT) and the other as overweight (beginning to slightly diverge from WT at 17 weeks, reaching 35% in extra-weight at 29 weeks) (Fig.1D). Intriguingly, although both fertility and weight parameters allowed us to identify subgroups of affected *Gw* mice in similar proportions (41% subfertile *vs* 40% overweight), these attributes were not strictly associated. Inclusion of all *Gw* mice that were evaluated for both parameters (also considering estrous cyclicity and induced ovulation data as additional proxies for fertility; both described in the next section) showed that only 26% (24/115) of affected *Gw* mice exhibited the combination of subfertility and overweight (Fig.1E). Importantly, transfer of the *Gw* allele onto the C57BL/6N genetic background, which is more resistant to metabolic dysfunction than FVB/N ^57^ and C57BL/6J ^58^ backgrounds, allowed the complete segregation of subfertility and overweight (Fig.S2). On the C57BL/6N background, 40% of *Gw* females still displayed subfertility (Fig.S2A), but in the complete absence of overweight (Fig.S2B). Thus, despite being known to negatively influence fertility ^31, 59^, overweight cannot be considered the cause of subfertility in *Gw* females. The dual association of subfertility and overweight with the *Gw* mutation in the FVB/N background, reminiscent of the metabolic subtype of PCOS, prompted us to further characterize both aspects of this model in more detail.

### Global alteration of reproductive functions in *Gw* females

Oligo/anovulation is a cardinal feature of PCOS, which we here examined by comparing estrous cyclicity in *Gw* and WT female mice at 2, 4, and 6 months of age. Based on the premise that overweight might aggravate reproductive deficits, we also analyzed lean and overweight subgroups of *Gw* mice independently (using weight at 6 months, implying retrospective assignment for earlier stages). Using vaginal smears to determine the number of estrous cycles over 16 days, we found that both lean and overweight *Gw* mice had fewer cycles than WT mice, a defect that worsened with age and was more pronounced in the overweight subgroup (Fig.3A). Closer examination of each phase of the estrous cycle at 6 months further revealed that both lean and overweight *Gw* mice spent more time in diestrus compared to WT mice, at the expense of both proestrus and estrus (Fig.3B). A similar trend was also noted at 4 months, but not earlier at 2 months (Fig.S3A). Consistent with this proxy of ovulatory dysfunction, the ovarian histology of both lean and overweight *Gw* mice revealed fewer corpora lutea and more abundant atretic follicles and follicular cysts compared to WT mice, at all three time points examined (Figs.3C-D and S3B). Moreover, we discovered that both lean and overweight *Gw* mice were less responsive than WT mice to artificially induced ovulation via PMSG and hCG injections at 4 and 6 months of age, with overweight further negatively impacting the quality of retrieved oocytes (Fig.3E-F).

**Figure 3.**
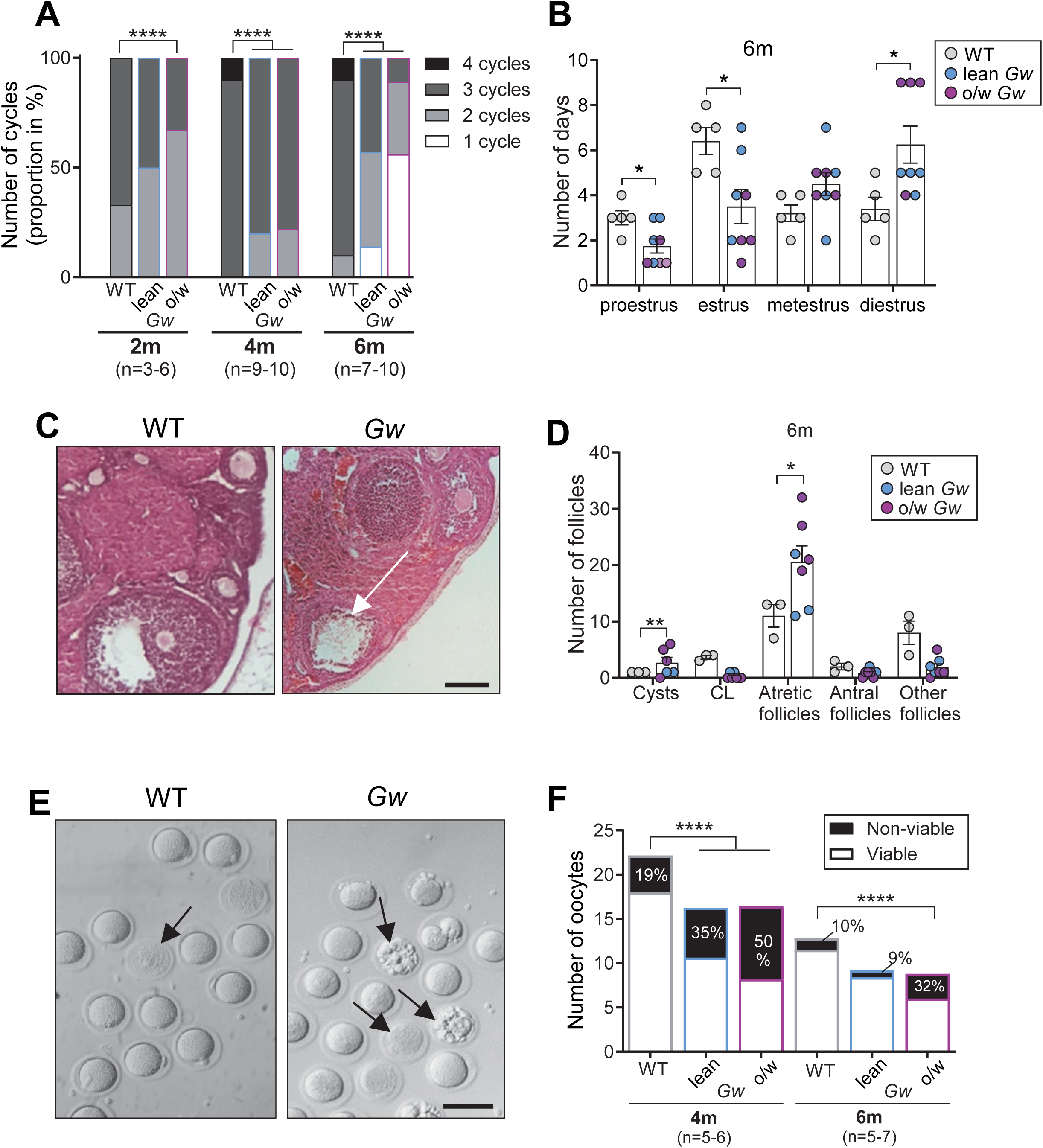
Estrous cycle profile, ovarian histology, and oocyte quality of *Gw* females are consistent with a PCOS-like phenotype. **A**) Number of estrous cycles per period of 16 days in WT and both lean and overweight (o/w) *Gw* female mice at 2, 4 and 6 months (m) of age (n=3-10 females per group). **B)** Number of days spent in each phase of the estrous cycle per period of 16 days in WT and both lean and overweight (o/w) *Gw* female mice at 6 months (m) of age (n=4-5 females per group). **C)** Hematoxylin & eosin staining of ovarian sections from 6-month-old ovaries showing a representative follicular cyst (white arrow) in *Gw* female mice. **D)** Quantitative analysis of the number of cysts, *corpora lutea* (CL), atretic follicles, antral follicles and other follicles counted on ovarian sections of 6-month-old WT and both lean and overweight (o/w) *Gw* female mice, using images such as those displayed in panel C. **E)** Gross morphology of oocytes collected from 4-month-old WT and *Gw* female mice, after artificially induced ovulation using PMSG and hCG. Arrows point to non-viable oocytes characterized by blurred or fragmented appearance. **F)** Quantification of the number and quality of retrieved oocytes after induction of ovulation, using images such as those displayed in panel E (n=5-7 induced females per group). Scale bar, 200 µm (C) and 50 µm (E). \**p* < 0.05, ** *p* < 0.01, **** *p* < 0.0001; chi-square tests (A and F) or two-tailed Welch’s *t*-test (B and D).

Using blood samples collected during the metestrus phase at 2, 4, and 6 months of age, we then examined plasma testosterone by ELISA. Average levels of testosterone in *Gw* mice were higher than in WT controls at all time points examined, regardless of whether they belonged to the lean or overweight subgroups (Fig.4A). Similar increases were detected for estradiol and LH (Fig.4A), as expected for these additional PCOS-relevant reproductive hormones. The impact of partial penetrance of the phenotype was especially evident for all hormonal measurements, leading to a high degree of data heterogeneity that prevented statistical significance from being reached at some time points. Like all three hormones mentioned above, higher levels of AMH are generally expected in a PCOS context ^60^, but showed no changes in *Gw* mice (Fig.S4A). As expected ^60^, no changes were noted for FSH levels (Fig.S4B). For LH, we additionally evaluated pulses in tail tip blood samples from 4-month-old mice, which revealed modestly, but significantly increased pulse frequency in *Gw* mice compared to WT controls (Fig.4B-C).

**Figure 4.**
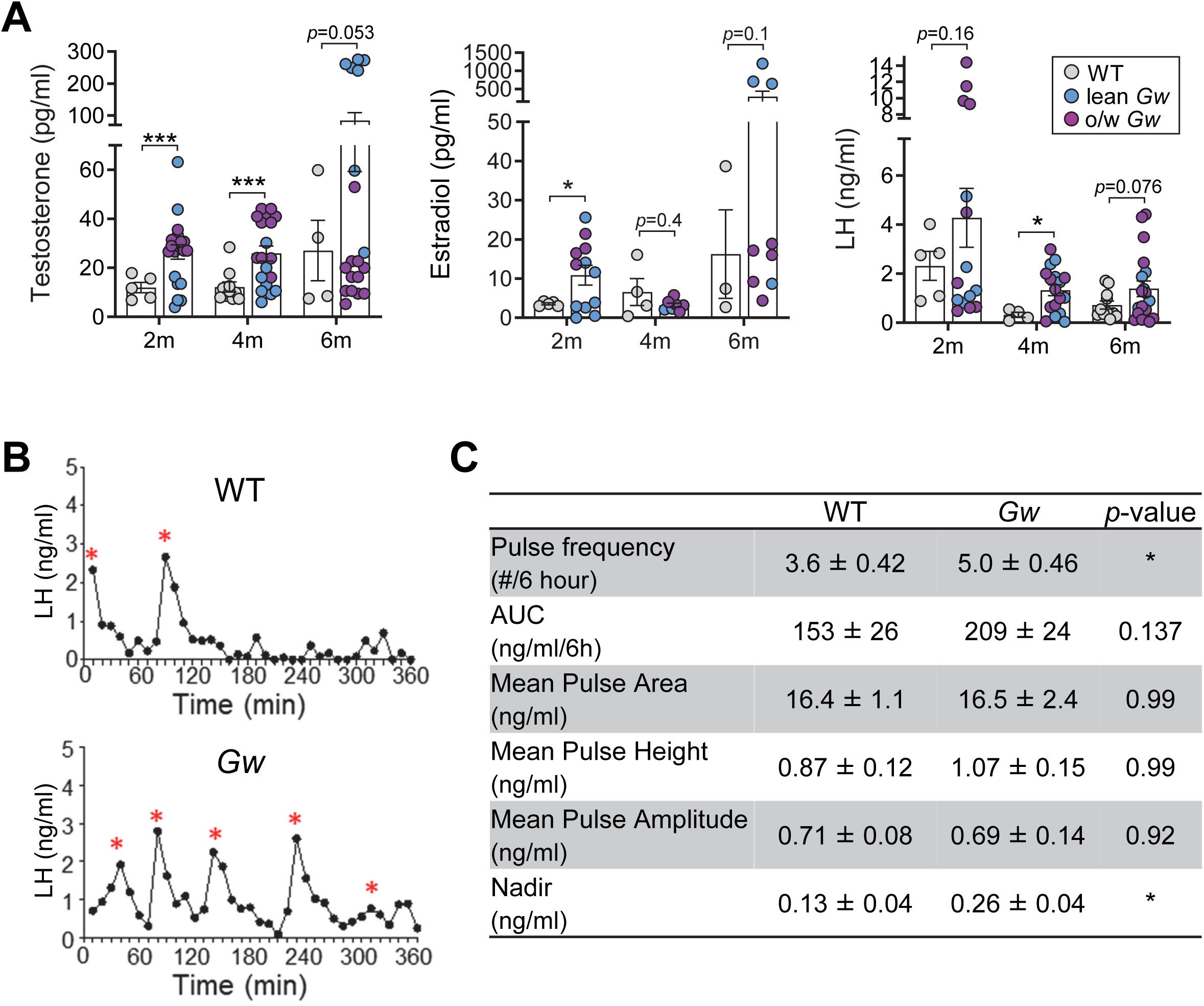
Hormonal profiles of *Gw* females are consistent with a PCOS-like phenotype. **A**) Circulating plasma levels of testosterone (left panel), estradiol (middle panel) and LH (right panel) in WT and both lean and overweight (o/w) *Gw* female mice during metestrus at 2, 4 and 6 months (m) of age (n=3-11 mice per group). **B)** Representative graphs of pulsatile LH levels in tail-tip blood samples taken every 10 min in 4-month-old female WT and *Gw* mice. Red asterisks indicate pulses. **C)** Statistical analysis of LH pulsatility using graphs such as those displayed in panel B (n=8 mice per group). \**p* < 0.05, *** *p* < 0.001; two-tailed Welch’s *t*-test.

We next examined *Gw* females during puberty, a stage of reproductive development critical for PCOS pathogenesis ^61^. Both the age of vaginal opening and the interval between the appearance of first estrus and the establishment of regular cycles were delayed in *Gw* females compared to WT controls (Fig.S5A-C). Delayed sexual development and maturation has been previously associated with low body weight ^62^, but this was not the case for *Gw* mice (Fig.S5D). Furthermore, although PCOS has been associated with longer anogenital distance (AGD), a biomarker of intrauterine fetal exposure to androgen ^63^, juvenile *Gw* females had significantly shortened AGD compared to WT control mice (Fig.S5E). This indicates that PCOS-like phenotypes in *Gw* females are not likely due to prenatal exposure to an excess of androgens, which is a common way to model PCOS in mice ^12, 13, 14, 15^. In sum, our data suggest that subfertility in *Gw* mice results from complex multi-layer perturbation of reproductive functions after birth.

### *Gw* mice exhibit a wide range of metabolic anomalies

To complement our analysis of weight gain in *Gw* females, we examined fat distribution and adipocyte size. At 4 months of age, we found that adipose tissue mass was greater in *Gw* females than controls at all sites that were examined, including perirenal, gonadal (paraovarian) and subcutaneous fat (Fig.S6A-D). At 6 months of age, a clear distinction could be made between overweight and lean subgroups of *Gw* mice, with this latter subgroup remaining nonetheless distinct from WT controls. For both subgroups of *Gw* females, we found that the increased fat mass could be explained by the presence of larger adipocytes, an observation especially striking in paraovarian fat pads (Fig.5A-B). Food consumption over a period of 4 days was greater in both subgroups of *Gw* mice compared to WT controls at this age (Fig.5C). Increased food intake by *Gw* mice was also confirmed at 4 months of age via automated measurements in metabolic cages (Fig.S6E). In line with this overfeeding phenotype, additional metabolic cage analyses revealed a higher respiratory quotient (RQ) in *Gw* mice (Fig.S6F), while ambulatory activity (XAMB) and energy expenditure (EE) were unaffected (Fig.S6G-H). Increased fat mass and overfeeding in *Gw* mice were, however, not found to be associated with a significant increase in plasma leptin levels measured by ELISA (Fig.5D).

**Figure 5.**
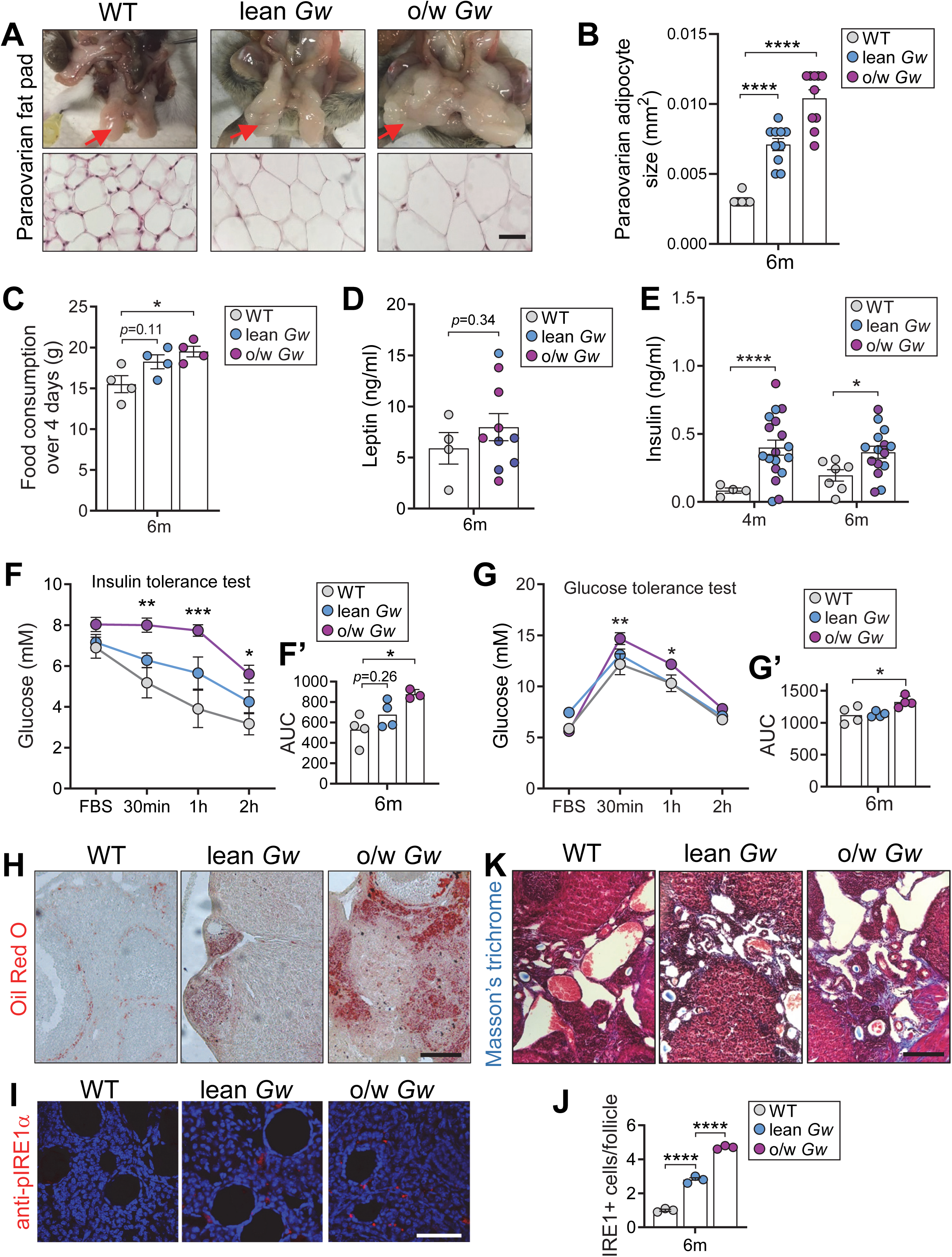
Metabolic parameters of *Gw* females are consistent with the metabolic subtype of PCOS. **A**) Relative size of paraovarian fat pads (red arrows, upper panels) and their H&E-stained adipocytes (lower panels) in WT and both lean and overweight (o/w) *Gw* female mice at 6 months of age (representative images of n=3 per group). **B)** Quantitative analysis of paraovarian adipocyte size in in 6-month-old mice, using images such as those displayed in lower panels in A (n=3 mice per group, average for 3 fields of view per sample). **C)** Food consumption of 6-month-old female mice over a period of 4 days. **D-E)** Circulating levels of leptin (at 6 months; D) and insulin (at 4 and 6 months; E) in adult female mice after 4h of fasting. **F-F’)** Glucose levels in 6-month-old female mice after 4h of fasting (FBS), and at indicated time points following i.p. injection of a 0.5 U/ml insulin solution (0.5 mU/g of body weight; n=3-4 mice per group). Data based on area under the curve (AUC) are shown in F’. **G-G’)** Glucose levels in 6-month-old female mice after 6h of fasting (FBS), and at indicated time points following i.p. injection of a 0.1 g/ml glucose solution (1 mg/g of body weight; n=4 mice per group). Data based on area under the curve (AUC) are shown in G’. **H, K)** Oil red O staining of neutral lipids (H) and Masson’s trichrome staining of collagen (K) in cross-sections of ovaries from 6-month-old female mice (representative images of n=3 mice per group). **I)** Immunofluorescence staining of the ER stress marker IRE1 (red) in cross-sections of ovaries from 6-month-old female mice, with cell nuclei counterstained using DAPI (blue). **J)** Quantitative analysis of the number of IRE1-positive cells in ovarian follicles using images such as those displayed in panel I (n=3 mice per group, average for 5 fields of view per sample). Scale bar, 100 µm (A), 200 µm (H) and 50 µm (I). * *p* < 0.05, ** *p* < 0.01, **** *p* < 0.0001; one-way ANOVA with Tukey’s post hoc test (B, C, F’, G’), two-tailed Welch’s *t*-tests (D and E), or two-way ANOVA with Tukey’s post hoc test (F and G).

We next examined whether the presence of enlarged adipocytes in *Gw* mice was associated with hyperinsulinemia and insulin resistance, as observed in women with PCOS ^64^. ELISA-based assessment of fasting plasma insulin levels at 4 and 6 months of age demonstrated hyperinsulinemia in both lean and overweight subgroups of *Gw* mice (Fig.5E). Yet, in the insulin tolerance test, only *Gw* females of the overweight subgroup showed impaired blood glucose uptake in response to an insulin injection, indicating insulin resistance (Fig.5F-F’). A similar outcome, although less marked, was observed in the glucose tolerance test, revealing that overweight *Gw* mice were also less able to efficiently manage a glucose injection (Fig.5G-G’).

We also tested whether the increased adiposity observed in *Gw* mice might impact ovaries locally, thereby potentially explaining why some reproductive parameters were more perturbed in the overweight subgroup (Fig.3). Enlarged adipocytes, as seen in *Gw* mice (Fig.5A-B), could reflect a triglyceride overload that can redirect the accumulation of fatty acids in ectopic locations ^65, 66^. We thus examined fat accumulation in *Gw* ovaries via Oil Red O staining of neutral lipids in ovary cross-sections from 6-month-old mice. While this dye only stained theca cells in control ovaries, some regions of ovarian cortex were also stained in *Gw* mice, and red staining was more intense in the overweight subgroup (Fig.5H). Excessive ectopic fat accumulation can sequentially lead to chronic low-grade inflammation and endoplasmic reticulum (ER) stress ^67^, which also contributes to ovarian fibrosis in the context of PCOS ^30, 68, 69^. We thus tested if this series of events was activated in *Gw* ovaries, via immunostaining of the ER stress sensor phospho-IRE1α (Fig.5I-J) and Masson’s trichrome staining of fibrosis-associated collagen (Fig.5K). All *Gw* samples displayed more numerous phospho-IRE1α-positive cells (within and around follicles) and more intense collagen staining (especially visible in the ovarian medulla), and these defects again looked more severe in the overweight subgroup. Taken together, our metabolic data thus suggest the existence of a pathogenic cascade in *Gw* mice that is likely triggered by impaired regulation of food consumption and eventually culminates in local negative impacts in ovaries.

### Downregulation of endogenous *Gata4* in the hypothalamus of *Gw* mice

Detailed characterization of the *Gw* insertion site (Fig.1F) and independent deletion of the associated *Gm10800* pseudogene (Figs.2 and S1) strongly suggested that *Gw* phenotypes are due to competition between transgenic and endogenous *Gata4* promoters, and hence downregulation of *Gata4* expression in a PCOS-relevant tissue. One such tissue is the ovary, where the GATA4 protein is notably present in both granulosa and theca cells of growing follicles ^40^. However, we knew from our prior work that gonadal expression of the *Gata4p[5kb]-RFP* transgene in the *Gw* line was restricted to testicular Sertoli cells, without any expression in ovaries ^44^. To verify the promoter competition hypothesis, we thus turned to another key tissue for PCOS pathogenesis, the hypothalamus. During murine hypothalamic development, the GATA4 protein is particularly abundant in migratory GnRH neurons ^70^, being also detectable in the immortalized GnRH neuron-like cell line, GT1-7 ^71, 72^. About a quarter of GnRH neurons are derived from neural crest cells ^73^, where the *Gata4p[5kb]-RFP* transgene is robustly expressed during prenatal development ^47^. Analysis of RFP emission in unfixed flat-mount preparations of hypothalamus from 4-month-old *Gw* mice (Fig.6) revealed that this transgene is apparently still active in postnatal GnRH neurons (recognizable by their thick dendron, with a high density of dendritic spines in the proximal region ^74, 75^), as we previously observed in other neural crest derivatives within the postnatal peripheral nervous system ^76^. Of note, this analysis further suggested that the transgenic *Gata4* reporter is active not only in GnRH neurons (Fig.6A) that are grouped in the anterior hypothalamus (combining preoptic area (POA) and anterior hypothalamic area (AHA)), but also in star-shaped astrocytes (Fig.6B) that are more widely scattered in the mediobasal hypothalamus (combining arcuate nucleus (Arc) and ventromedial hypothalamus (VMH)). This latter observation is consistent with prior studies reporting the presence of endogenous GATA4 protein in postnatal astrocytes from other brain regions of both mice and humans ^77, 78, 79^, and we confirmed by RT-qPCR that *Rfp* transcripts are present in Arc and VMH collected separately from 4-month-old *Gw* mice (Fig.6C).

**Figure 6.**
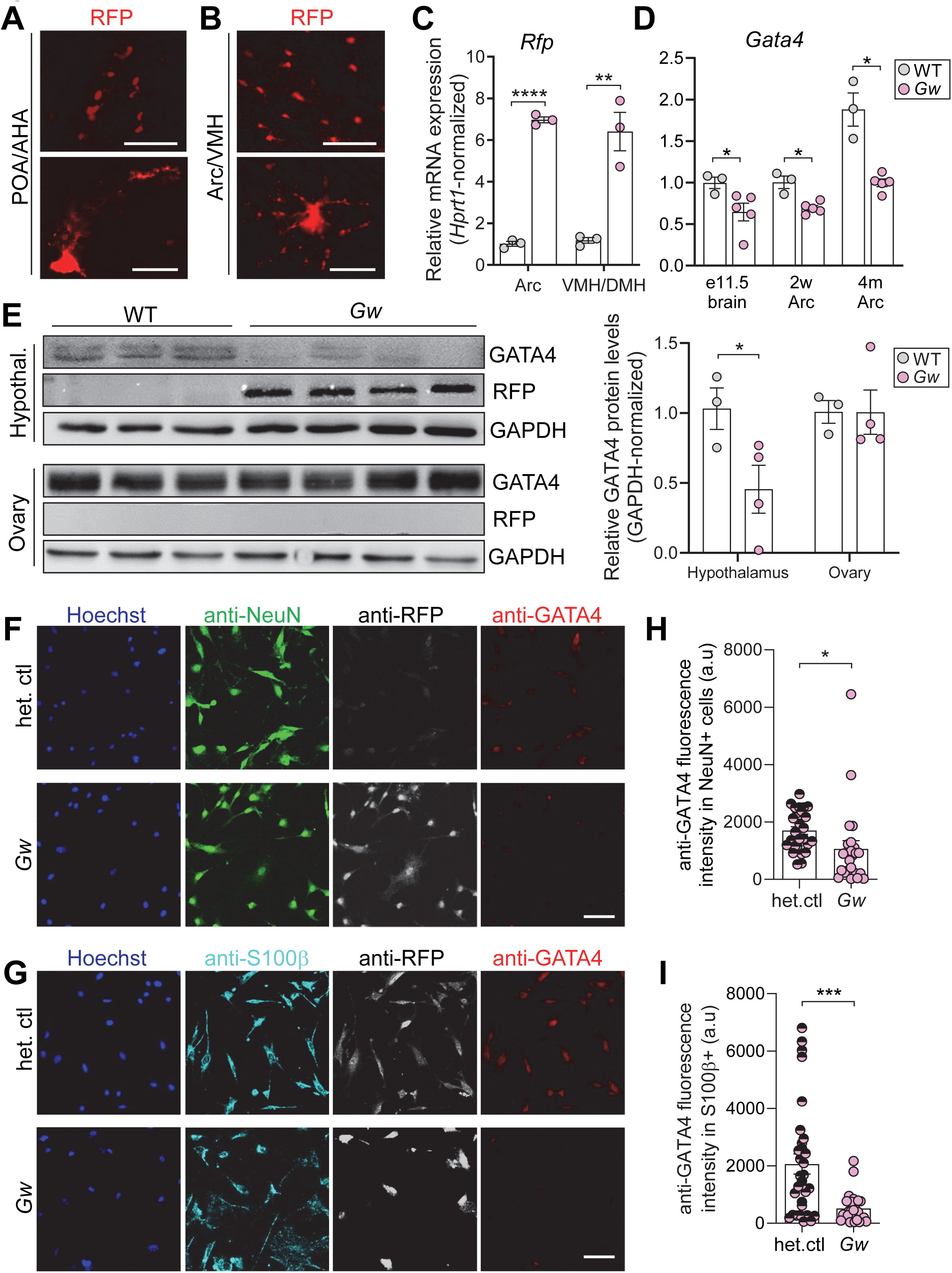
Expression of the *Gata4p[5kb]-RFP* transgene in the hypothalamus of *Gw* female mice is associated with downregulation of endogenous *Gata4*. **A**) Distribution (upper panel; low magnification) and morphology (lower panel; high magnification) of RFP-positive cells in preoptic (POA) and anterior (AHA) hypothalamus areas of unfixed/flatmounted hypothalamus from 4-month-old *Gw* female mice is consistent with a GnRH neuron identity (representative of n=3 mice). **B)** Distribution (upper panel; low magnification) and morphology (lower panel; high magnification) of RFP-positive cells in arcuate nucleus (Arc) and ventromedial hypothalamus (VMH) of unfixed/flatmounted hypothalamus from 4-month-old *Gw* female mice is consistent with an astrocyte identity (representative of n=3 mice). **C)** RT-qPCR analysis of *Rfp* expression in 4-month-old Arc and VMH (also including dorsomedial hypothalamus, DMH) nuclei, microdissected after slicing in brain matrix (n=3 mice per group). **D)** RT-qPCR analysis of *Gata4* expression in developing brain (e11.5) as well as pubertal (2 weeks) and adult (4 months) Arc nucleus, microdissected after slicing in brain matrix (n=3-5 mice per group). **E)** Western blot analysis of GATA4 protein levels in whole hypothalamus and ovary from 4-month-old WT and *Gw* female mice (n=3-4 mice per group), with accompanying quantification shown on the right. **F)** Representative confocal images of NeuN+ hypothalamic neurons (green) co-stained for Hoechst (blue), RFP (white) and GATA4 (red) after dissociation from 1-month-old heterozygous (het. ctl) and homozygous *Gw* hypothalamus and 7 days of culture (n=3 mice per group). **G)** Representative confocal images of S100β+ hypothalamic astrocytes (green) co-stained for Hoechst (blue), RFP (white) and GATA4 (red) after dissociation from 1-month-old heterozygous (het. ctl) and homozygous *Gw* hypothalamus and 7 days of culture (n=3 mice per group). **H-I)** Quantification of GATA4 protein levels in cultured hypothalamic cells based on intensity of immunofluorescence staining, using images such as those displayed in panels F-G (n=3 mice per group, 3-5 fields of view per sample). Scale bar, 100 μm (upper panels in A-B), 25μm (lower panels in A-B) and 50 μm (F-G). * *p* < 0.05, ** *p* < 0.01, **** *p* < 0.0001; two-tailed Welch’s *t*-tests.

In accordance with the promoter competition hypothesis, we found that endogenous *Gata4* mRNA expression was decreased by about 50% in the Arc of 4-month-old *Gw* mice, while smaller decreases were observed in juvenile Arc and embryonic brain (Fig.6D). The specificity of *Gata4p[5kb]-RFP* transgene expression and *Gata4* downregulation in 4-month-old *Gw* mice was further validated by western blotting, which we also use to confirm that GATA4 protein levels were not affected in RFP-negative ovaries (Fig.6E). To directly verify the impact of *Gata4p[5kb]-RFP* transgene activity on endogenous *Gata4* expression, we next sought to isolate RFP+ cells from the hypothalamus of *Gw* mice by FACS (fluorescence-activated cell sorting) for more targeted RT-qPCR analyses – using heterozygous *Gw* mice as controls with ∼50% lower transgene expression levels (Fig.S7). However, we repeatedly failed to isolate high-quality RNA in sufficient amount for these experiments, most likely because of a combination of low cell numbers (<1%; Table S1) and sensitivity to tissue dissociation and/or flow cytometry. To circumvent this problem, we cultured freshly dissociated hypothalamic cells for 7 days in a medium that normally supports the growth and survival of both neurons and astrocytes, thereby allowing their analysis by immunofluorescence. These experimental conditions also offered the advantage of making our GATA4 and RFP antibodies now usable for immunostaining, which otherwise did not work for us on either sections or whole-mount preparations of the hypothalamus. Importantly, comparative analysis of cultured hypothalamic cells from *Gw* mice and their heterozygous controls at 1-month of age revealed a reduction of GATA4 immunoreactivity in both NeuN-positive neurons (Fig.6F,H) and S100β-positive astrocytes (Fig.6G,I) from *Gw* mice, and this downregulation was systematically accompanied by an increase in RFP immunoreactivity (Fig.6F-G). We also noted that neurons and astrocytes mostly grew independently of each other, suggesting that our culture conditions favored clonal expansion and differentiation from pre-committed tissue-resident progenitors – as expected from a neurogenic/gliogenic center like the hypothalamus ^80, 81, 82^ – rather than maintenance of pre-existing neurons and astrocytes. Hence, the scarce GnRH neurons, which originate from extra-hypothalamic progenitors, were most likely excluded from this analysis. Nevertheless, regardless of the exact identity of *Gata4*-expressing neurons and glia that are present in these experiments, our results strongly support the promoter competition hypothesis.

### Perturbed gene expression related to hypothalamic regulation of reproduction and food intake in *Gw* mice

GATA4 being a transcription factor, we reasoned that its decreased levels in the hypothalamus of *Gw* mice would perturb – either directly or indirectly – the gene expression program that controls reproductive functions and food intake. For these studies, we specifically focused on the Arc from 4-month-old mice (where and when *Gata4* is clearly downregulated in *Gw* mice; Fig.6E). Using RT-qPCR, we first analyzed transcript levels of the mediators of ovarian hormone negative feedback that are expressed in various populations of neurons and/or astrocytes. This analysis revealed a significant downregulation of the estrogen receptor-coding gene *Esr1* in *Gw* samples compared to WT (Fig.7A), with immunostaining further revealing that this decrease is not due to a lower number of ERα-positive cells in the Arc (Fig.S8A-B). A trend toward reduced expression of *Pgr* was also noted in *Gw* samples, but without reaching statistical significance (Fig.7A). As neuronal proteins under external stimuli regulation in dendrites/axons are often translated locally ^83, 84^, we also analyzed transcript levels of the main GnRH isoform (*Gnrh1*) in projections from POA-located GnRH neurons passing through the Arc (where a few cell bodies of GnRH neurons are present as well ^85, 86^). As observed for *Pgr*, no significant difference was found for *Gnrh1* expression levels (Fig.7A). Yet, we observed opposite trends for *Gnrh1* transcript levels (slightly increased in *Gw*) and the number of GnRH neurons in the POA (slightly decreased in *Gw*) (Fig.S8C-D), suggestive of a potential compensatory mechanism acting at the transcription level. In addition, the analysis of key regulators of pulsatile GnRH release showed a significant upregulation of the positive regulator *Kiss1* in *Gw* samples, with no significant difference noted for the other positive regulator *Nknb* and the negative regulator *Pdyn* (Fig.7B). Lastly, analysis of regulators of food intake revealed a significant downregulation of the anorexigenic *Pomc* in *Gw* samples, with no significant difference noted for the orexigenic *Agrp* and *Npy* (Fig.7C). This was accompanied by a specific increase in mRNA levels of the short (non-signaling) isoform of the leptin receptor, without significant impact on mRNA levels of the long (signaling) isoform (Fig.7C). All these results provide a mechanistic basis for eventually understanding the reproductive and metabolic phenotypes of *Gw* mice, suggesting that these mice might: i) be less sensitive to estrogen negative feedback, ii) have increased GnRH pulsatility, and iii) have impaired satiety.

**Figure 7.**
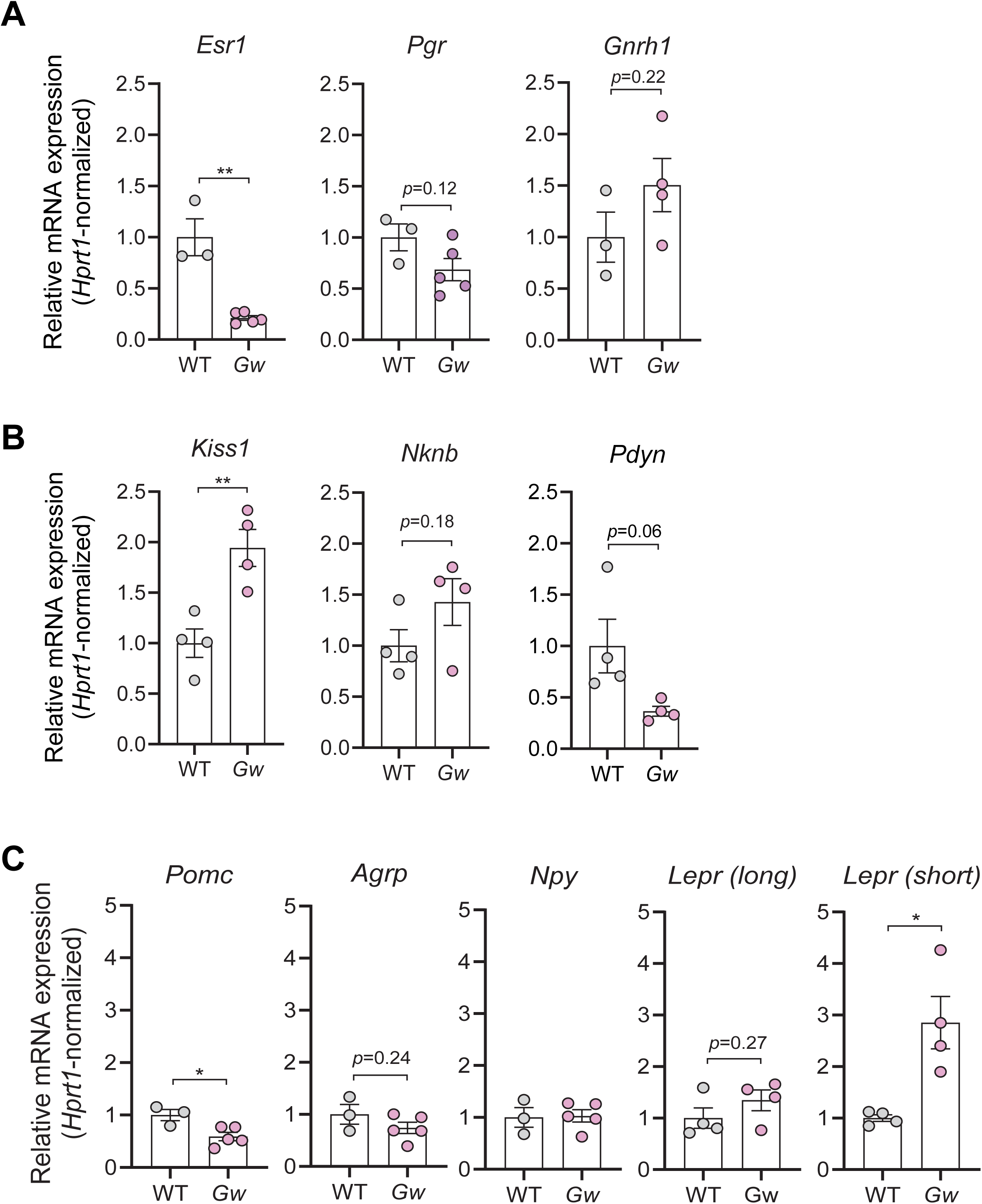
Multiple markers of hormonal feedback, GnRH release and food intake are dysregulated in the hypothalamus of *Gw* female mice. **A-C**) RT-qPCR analysis of sex hormone receptor-coding genes (A), GnRH pulse generator-coding genes (B), and food intake-associated neuropeptide-coding genes (C) in 4-month-old Arc nucleus, microdissected from WT and *Gw* mice after slicing in brain matrix (n=3-5 mice per group). * *p* < 0.05, ** *p* < 0.01; two-tailed Welch’s *t*-tests.

### Postnatal brain-specific knockout of *Gata4* in either neurons or astrocytes impairs fertility in female mice

In a final set of experiments, we independently examined the role of GATA4 in the postnatal brain while also investigating its relative contribution in neurons and astrocytes. To this end, we generated female mice homozygous for the conditional *Gata^LoxP^* allele ^91^ also bearing a tamoxifen-inducible Cre driver transgene active in neural stem cells and subsets of either neurons (using *Nestin-CreERT2* ^92^) or astrocytes (using *GFAP-CreERT2* ^93^). We then evaluated fertility rates and weight gain in these *Gata4^LoxP/LoxP^;Nestin-CreERT2^Tg/+^*and *Gata4^LoxP/LoxP^;GFAP-CreERT2^Tg/+^* females (all in FVB/N background) following tamoxifen injections at 1 month of age, using Cre-negative littermates as controls (Fig.8A). For each conditional knockout line, the expected partial decrease of total hypothalamic GATA4 protein levels was confirmed by western blotting (Fig.8B), while the cell type specificity of Cre activation in either neurons or astrocytes was independently confirmed using the Cre reporter allele *R26^[Floxed^ ^Stop]YFP^* ^94^ (Fig.8C). Remarkably, we found lower fertility rates in about half of tamoxifen-treated female mice of both conditional knockout models between 2 to 6 months of age (Fig.8D), as previously observed in *Gw* mice (Fig.1B). However, neither conditional approach appeared to significantly impact body weight at 4 months (Fig.8E). Interestingly, cell lineage-specific effects were noted when analyzing pregnancy complication-related parameters. Indeed, while high rates of miscarriage were observed in both conditional knockout models, an abnormally high rate of perinatal death was only present in *Gata4^LoxP/LoxP^;GFAP-CreERT2^Tg/+^*females (Fig.8F). Altogether, these data validate that GATA4 is needed in the brain for the regulation of three PCOS-relevant fertility/pregnancy parameters that are affected in *Gw* mice, with some regulatory aspects being either partly shared by neuronal and astrocytic GATA4 (fertility/miscarriage rates) or exclusive to astrocytic GATA4 (perinatal death rate). These data also suggest that the extra-weight gain seen in *Gw* mice might be due to the combined decrease of GATA4 in both neurons and astrocytes.

**Figure 8.**
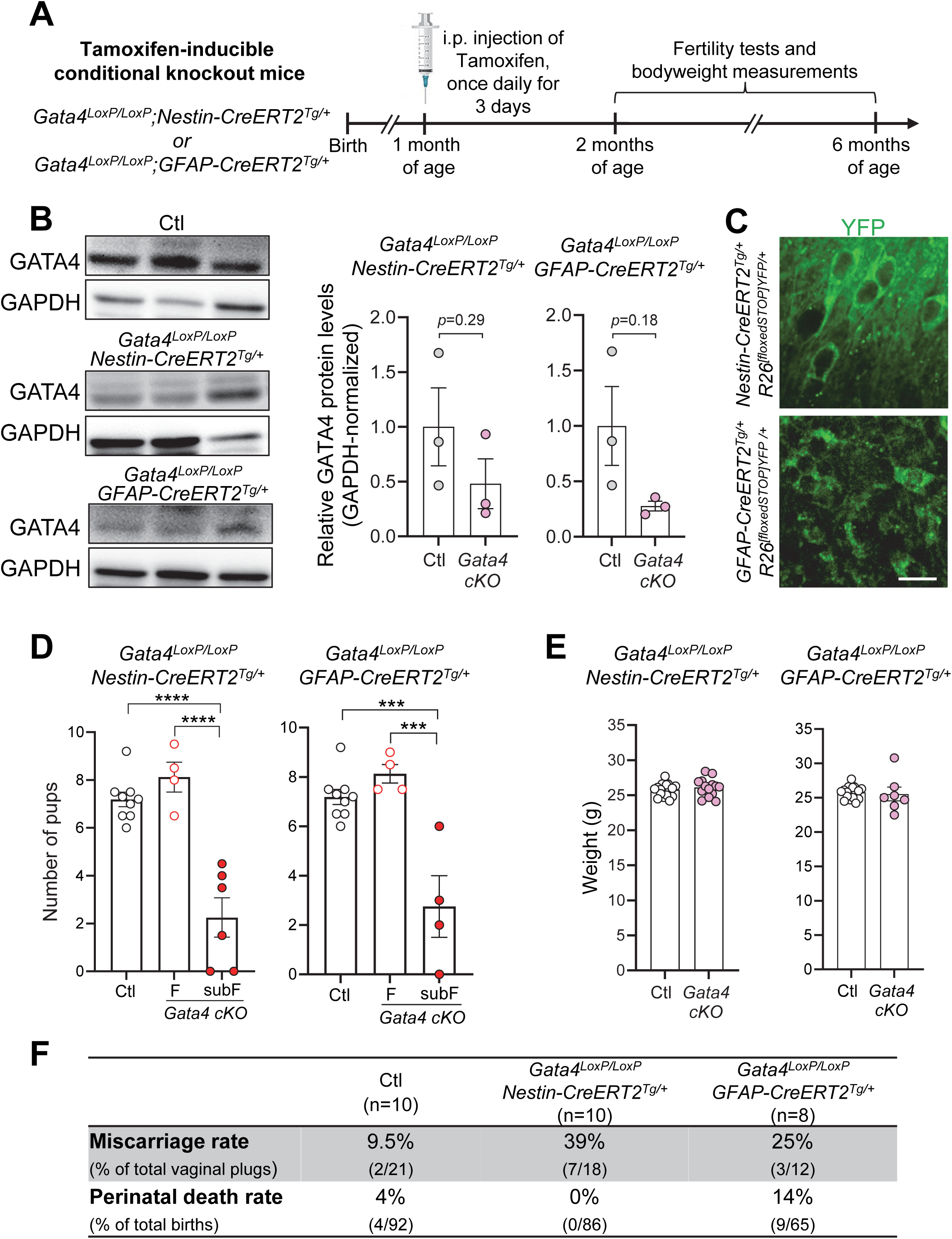
Brain-specific knockout of *Gata4* in either neurons or astrocytes phenocopies the subfertility phenotype of *Gw* female mice. **A**) Overview of conditional knockout strategies and experimental design for assessment of fertility and bodyweight in resulting tamoxifen-treated *Gata4^LoxP/LoxP^;Nestin-CreERT2^Tg/+^*and *Gata4^LoxP/LoxP^;GFAP-CreERT2^Tg/+^* female mice. **B)** Western blot analysis of GATA4 protein levels in whole hypothalamus of 4-month-old control and tamoxifen-treated *Gata4^LoxP/LoxP^;Nestin-CreERT2^Tg/+^* and *Gata4^LoxP/LoxP^;GFAP-CreERT2^Tg/+^* female mice (n=3 mice per group), with accompanying quantification shown on the right. **C)** Validation of tamoxifen-induced Cre activity in neurons (left panel) and astrocytes (right panel), based on *Rosa26* locus-driven YFP expression in tamoxifen-treated *Nestin-CreERT2^Tg/+^;Rosa26^[FloxedSTOP]YFP/+^* and *GFAP-CreERT2^Tg/+^;Rosa26^[FloxedSTOP]YFP/+^* female mice (1 week after tamoxifen administration at 1 month of age), respectively. **D)** Average number of pups per litter for control (same group in both graphs) and tamoxifen-treated conditional *Gata4* knockout females (2-to 4-month-old), after breeding with WT males (n=4-9 females per group, 2-3 litters per female). F, fertile; subF, subfertile. **E)** Bodyweight of 4-month-old control (same group in both graphs) and tamoxifen-treated conditional *Gata4* knockout female mice before breeding (n=7-21 females per group). Note that there is a single control group, graphed twice for presentation purposes. **F)** Comparative table of pregnancy complication-related parameters. Scale bar, 50µm. * *p* < 0.05, *** *p* < 0.001; two-tailed Welch’s *t*-tests.

## DISCUSSION

Initially triggered by a serendipitous discovery in the *Gw* transgenic mouse line, this study finally provides a solid foundation to understand the association between *GATA4* SNPs and PCOS. When also considering the analysis of conditional knockout mouse lines, our work strongly suggests that this link with PCOS is due to perturbation of a previously overlooked role of GATA4 in the hypothalamus. Hence, another important aspect of this study is that it strengthens the notion that hypothalamic dysfunction can be at the root of PCOS.

### Usefulness of *Gw* mice for exploring pathogenic mechanisms and therapeutic avenues in PCOS

Adult *Gw* female mice will continue to be useful for future studies aimed at clarifying PCOS pathogenesis, with or without overweight. Even the partial penetrance of PCOS-associated phenotypes in this mouse line is of interest, as it mimics the heterogeneous presentation observed in human cases^3^. The main advantage of the *Gw* mouse model is that PCOS-like traits develop spontaneously, although it takes some time for the disease to clearly manifest itself (≥4 months). PCOS-like pathology generally develops more rapidly in hormone-induced models, but phenotypic presentation can vary considerably depending on hormone treatment regimen and both time and duration of administration ^13, 14, 15^. For instance, in androgen-induced models, pre-natal administration delays the presentation of metabolic phenotypes by several weeks whereas neonatal administration totally fails to induce PCOS-like features. These limitations are not a problem with later administration of androgens at puberty or adulthood, but the combination of resulting phenotypes again varies as a function of experimental conditions – most especially duration of treatment, which can last from a few weeks to an uninterrupted period. It is also important not to forget that environmental factors, including chow diet used in the animal facility and stress induced by the experimenter, can greatly impact phenotypic presentation as well ^14^. In the end, the choice of androgen treatment paradigm is often dictated by the specific aspects of PCOS one wishes to study, which is nonetheless greatly facilitated by the generally high penetrance of androgen-induced phenotypes. The letrozole-induced model has been reported to outperform the androgen-induced models in terms of co-occurring PCOS-related phenotypes ^95^. However, here again, there are some treatment regimen-dependent variability issues attached to this modeling approach in both mice ^96^ and rats ^97^. Moreover, the aromatase inhibitor action of letrozole (*i.e.*, preventing testosterone to estrogen conversion) is used in the clinic to treat infertility in women with PCOS ^98, 99^, complicating its use for disease modeling.

The spontaneous development of PCOS-like traits in *Gw* mice also offers a key advantage for therapeutic investigations. Indeed, artificial hormone-induced models can be useful for developing specific countermeasures, but much less for more global interventions that would ideally prevent the same hormonal problems that are used for PCOS modeling. To the best of our knowledge, spontaneous development of PCOS-like phenotypes has been reported in only one other rodent model, the Goto-Kakizaki (*GK*) rat ^100^. This rat line is best known as a model for non-obese type 2 diabetes mellitus, resulting from an artificial selection scheme for glucose intolerance over several generations ^101^. In terms of PCOS modeling, the *GK* rat line mainly differs from the *Gw* mouse line in genetics (polygenic for *GK vs.* transgenic for *Gw*), metabolic phenotype (lean only for *GK vs.* both lean and overweight for *Gw*) and life stage of phenotype presentation (puberty for *GK vs.* adults for *Gw*). This complementarity between both models could be an asset for therapy development, opening the possibility to identify therapeutics with either wide or more restricted applications.

### A previously overlooked role for GATA4 in the hypothalamus

Our mouse genetic data indicate that competition between transgenic (owing to integration in a strong enhancer region) and endogenous promoters is the underlying mechanism for the specific downregulation of *Gata4* expression in the hypothalamus of *Gw* female mice. Gene and protein expression data further show that hypothalamic GATA4 levels in adult *Gw* females correspond to about 50% of those seen in WT mice, raising the question as to why PCOS-like phenotypes have not been previously reported for heterozygous *Gata4*-null mice. This is most likely due to partial compensation from the remaining functional allele, which allows heterozygous *Gata4*-null mice to have in fact ∼40% less *Gata4* transcripts (i.e., ∼60% of WT levels) ^102^. As lowering GATA4 protein levels by 70% (i.e., 30% of WT levels) is also known to be embryonic lethal ^103^, all these data highlight the fact that GATA4 is highly sensitive to dosage, with a threshold level for not developing disease between 50-60% of WT levels. The fact that GATA4 levels are around the lower limit of this threshold in *Gw* mice might be the main reason for the incomplete phenotypic penetrance.

Further work is needed to elucidate the apparently complex role played by GATA4 in the hypothalamus. In particular, we need to clarify the identity of *Gata4*-expressing cell types in the postnatal hypothalamus of WT mice. Our current data suggest that these cells are relatively rare (Table S1) and include both neurons and glia (Fig.6). The recently described *Gata4^CreERT^*^2^ mouse line ^104^ could be especially useful in this regard, allowing us to overcome our problems with GATA4 immunostaining of brain sections via tamoxifen-inducible genetic tracing in conjunction with a Cre reporter line like *R26^[Floxed^ ^Stop]YFP^* ^94^. A better understanding of the *Gata4* expression profile in the hypothalamus will then enable the use of more precise conditional knockout approaches than those described in the current study. This will also be essential for designing and interpreting studies aimed at identifying the direct target genes of GATA4 in the hypothalamus. As seen with many transcription factors ^105, 106^, these targets can be expected to be regulated not only at the transcriptional level ^107, 108^, but also by co-transcriptional alternative splicing ^109^.

In addition, further research should focus on the two human *GATA4* SNPs (rs804279 and rs3729853) that have been linked with PCOS in GWAS data ^34, 36, 38^. For this work, it would also be important to analyze the contribution of a third association signal identified by gene-based association testing (summing the effects of all variants within a gene) and assigned to *LINC02905* (also known as *C8orf49*) ^38^, a putative non-coding RNA located in the *GATA4-NEIL2* intergenic region at position Chr8:11,761,256-11,763,223. Indeed, upon careful examination in the UCSC genome browser (not found in the Ensembl browser), *LINC02905/C8orf49* has all the hallmarks of a distal regulatory element, with ChIP-seq data notably showing enrichment of the H3K27Ac histone mark and pointing to a transcription factor binding hotspot. Both *GATA4* SNPs associated with PCOS (rs804279 at position Chr8:11,623,889 and rs3729853 at position Chr8:11,756,807) are located in regions with a similar signature, suggesting that all three regions could thus act as distal downstream enhancers directing *GATA4* expression in hypothalamic cells. This exciting possibility has potentially important implications beyond *GATA4*-associated PCOS, as it might help to identify functional links with some of the other signaling molecules / transcription factors that have also been associated with PCOS in GWAS data ^3^.

## METHODS

### Mice

The generation of the *Gw* transgenic mouse line (*Gata4p[5kb]-RFP* line #2) has been previously described ^49^. Briefly, one-cell FVB/N embryos were injected with a linearized rat *Gata4* promoter [5kb] – dsRed2 fluorescent protein construct (G4-RFP) using standard pronuclear microinjection methods ^110^. A previously described *tyrosinase* minigene construct ^49^ was co-injected to allow visual identification of transgenic animals via rescue of pigmentation of the FVB/N albino background ^51^. Transfer of the *Gw* allele on the C57BL/6N background was achieved by consecutive backcrossing over 5 generations. Genotyping of adult *Gw* animals in the FVB/N background was made by visual inspection of coat color whereas genotyping of adult *Gw* animals in the C57BL/6N background was performed by standard PCR using primers listed in Table S2.

*Gm10800* and *Gm10800/Gm10801* knockout mice were generated in the FVB/N background by the Transgenesis and Genome Editing core of the CERMO-FC research center, using the Easi-CRISPR approach as previously described ^111^. All CRISPR reagents were purchased from Integrated DNA Technologies (IDT), with guide RNAs designed to target just outside each end of the *Gm10800* locus at position Chr2:98,666,137 and position Chr2:98,667,443 (based on GRCm38 genome assembly). Each custom crRNA containing the relevant *Gm10800*-targeting guide RNA (5’-GAAGGGGAAUUUUGGUGGCA or 5’-ACAUGUUCAAAAGGGGCCCG) was annealed with the universal tracrRNA (IDT cat.#1072534), and resulting crRNA/tracrRNA complexes (final concentration of 0.61µM for each) were then combined with the Cas9 protein (IDT cat.#1081058; final concentration of 0.3µM) to obtain a <u>c</u>rRNA/tracr<u>RN</u>A Cas9 <u>P</u>rotein (ctRNP) complex that was finally microinjected in FVB/N zygotes using standard pronuclear microinjection methods ^110^. Founder animals were identified by PCR (see genotyping primers in Table S2), with deletion then carefully mapped using Sanger sequencing. Two founders (one with single deletion of *Gm10800*, and one with double deletion of the juxtaposed *Gm10800/Gm10801*) were kept and bred with FVB/N animals to establish heterozygous lines, which were then separately intercrossed to homozygosity.

*Gata4^LoxP/+^* (*Gata4<tm1.1Sad>/J*; JAX stock #008194), *Nestin-CreERT2^Tg/+^* (*C57BL/6*-*Tg[Nes-cre/ERT2]KEisc/J*; JAX stock # 016261) and *GFAP-CreERT2^Tg/+^* (*B6.Cg-Tg[GFAP-cre/ERT2]505Fmv/J*; JAX stock #012849) mice were purchased from The Jackson Laboratory (Bar Harbor, USA). All three lines were backcrossed in the FVB/N background for 5 generations before intercrossing *Gata4^LoxP/+^* with either of the Cre driver lines to generate *Gata4^LoxP/LoxP^; Nestin-CreERT2^Tg/+^* and *Gata4^LoxP/LoxP^; GFAP-CreERT2^Tg/+^* mice. To induce deletion of *Gata4* exons 3-5 in the postnatal brain, these mice were administered tamoxifen (Sigma cat.#T5648; diluted in corn oil at 10mg/ml) via single daily intraperitoneal injections of 1mg per kg of bodyweight for 3 consecutive days at 1 month of age. Efficiency of Cre-mediated recombination using this experimental design was confirmed for each Cre driver line after breeding with *R26^[FloxedSTOP]YFP^* (*Gt[ROSA]26Sortm1(EYFP)Cos*) mice, provided by F. Costantini (Columbia University, USA) and maintained on an FVB/N background ^94^. All studies on tamoxifen-treated control and mutant mice were then carried out between the ages of 2 to 6 months.

All mice were maintained and bred in individually ventilated cages within the conventional animal facility of the Université du Québec à Montréal, with *ad libitum* access to Charles River Rodent *Diet* #*5075* (Cargill Animal *Nutrition*) under 12h light – 12h dark cycles (7AM to 7PM). All experiments were approved by the local ethics committee of Université du Québec à Montréal (CIPA # 931) and performed according to guidelines of the Canadian Council on Animal Care (CCAC). Mice were euthanized by either CO_2_ inhalation or decapitation (for collecting brain tissues), both after isoflurane anesthesia. For timed pregnancies, mice were mated overnight, and noon on the day of vaginal plug detection was designated as E0.5.

### Mapping of Greywick transgene insertion site

Whole genome sequencing of homozygous *Gw* mice was performed at the Génome Québec Innovation Centre on three separate occasions, using different technologies. The first attempt used the HiSeq 2000 platform (Illumina), yielding 251 million of 100pb-long paired-end reads. The second attempt used the HiSeq X platform (Illumina), yielding 479 million of 150pb-long paired-end reads. The third attempt used the linked-reads approach (10x Genomics) prior to sequencing on the Hiseq X platform (Illumina), yielding 389 million 150pb-long paired-end reads. In each case, sequences were mapped onto the Mus musculus reference genome (GRCm38 assembly), to which transgenic sequences (*Gata4p[5kb]-RFP* and *tyrosinase* minigene) were added. Using these transgenic sequences as viewpoint on the Integrative Genomics Viewer (IGV) allowed precise mapping of both transgene-genome boundaries, which were further validated using PCR (see relevant primers in Table S2) and Sanger sequencing.

### Reproductive phenotyping

To examine fertility rates, mice of control and experimental groups to be tested were naturally mated with fertile WT animals of opposite sex (all aged between 2 to 6 months). Vaginal plugs were checked every morning and coitus resulting in pregnancy was determined by weight gain at mid-gestation. The number of delivered pups was then counted for at least two pregnancies per tested animal. Miscarriage was concluded when pregnant females stopped gaining weight after mid-gestation and did not give birth around the expected delivery date. Perinatal death rates were established by counting the number of live and death pups at birth and over the next two days.

To monitor the different phases of the estrous cycle, vaginal epithelial cells were collected from WT and Gw females (2, 4 and 6 months of age) via daily lavage of the opening of the vaginal canal with saline over a 16-day period, as previously described ^112^. The vaginal fluid collected was placed on a clean glass slide and stained with crystal violet (0.1% solution). Stage of the estrous cycle was determined by examining smears under a Leica DM2000 histology microscope. The proestrus phase was identified by the presence of primarily nucleated epithelial cells. The estrus phase was identified by the presence of predominantly cornified epithelial cells. The diestrus phase was identified by the presence of predominantly leucocytes. The metestrus phase was identified by a mixture of nucleated cells, cornified cells, and leucocytes.

To evaluate oocyte quality, WT and Gw females (4 months of age) underwent superovulation by intraperitoneal (IP) injection of 5 IU pregnant mare’s serum gonadotrophin (PMSG; Sigma Cat.# G4877-1000 IU, followed 48 h later by injection of 5 IU of human chorionic gonadotrophin (hCG) (Sigma Cat.# CG10). Thirteen to fourteen hours after hCG injection, animals were sacrificed, and oocytes were collected from the fallopian tubes. Oocytes were finally examined under a Leica M80 stereomicroscope to determine their number, morphology and viability.

### Metabolic phenotyping

The body weight of WT and *Gw* female mice were assessed weekly using a standard laboratory balance (Ohaus) and then *Gw* mice were divided into lean and overweight groups. Fat pads from gonadal, subcutaneous, perirenal and pericardial were carefully dissected out from 4-month-old mice and weighed on an analytical balance (Shimadzu).

To globally assess functional metabolism and food consumption, WT and *Gw* females (4 months of age) were individually housed in metabolic cages equipped with the Comprehensive Laboratory Animal Monitoring System (CLAMS; Columbus Instruments). As recommended by the manufacturer, each mouse was supplied with enough food for the duration of data collection (three consecutive days). All mice were acclimatized for 24h before measurements. Analyses were made using data collected at 48h, which allows adequate time to acclimate to the CLAMS environment and provide accurate data for assessment of all measured parameters. O2 consumption (VO2), CO2 production (VCO2), Respiratory Exchange Rate (RER), food intake (FI), Energy Expenditure (EE), as well as X-Ambulatory counts (XAMB, physical activity) were determined. The RER was calculated as the ratio of carbon dioxide production and oxygen consumption. Energy expenditure was calculated using the following equation: heat = CV x VO2, where CV= 3.815+1.232 x respiratory exchange ratio (CV, calorific value based on the observed respiratory exchange ratio). XAMB locomotor activity per day was determined for each mouse by counting the movement made across the cage, which is measured with infrared sensors. Measurements of food intake with CLAMS were broken down into light and dark cycles for a total of 48h per mouse. In addition to CLAMS, food intake was manually measured for groups of four mice per cage by weighing the food every four days over a period of 30 days to calculate the average consumption of food/mouse/day in each group.

For glucose and insulin tolerance tests, WT and *Gw* females (4 and 6 months of age) were fasted for either 4h (8:00AM-12:00PM; for insulin tolerance test) or 6h (8:00AM-2:00PM; for glucose tolerance test). Blood glucose was determined in a drop of whole blood using a glucometer (Accu-Check Aviva, Roche). Blood glucose concentrations were measured at 30, 60, and 120 min after intraperitoneal injection of 1g/kg glucose (glucose tolerance test) or 0.5 U/kg of recombinant human insulin (Sigma Cat.# I6634) (insulin tolerance test).

### Hormonal analyses

Insulin and leptin levels were measured in plasma samples obtained from 6h fasted WT and *Gw* females (4 and/or 6 months of age) using ELISA kits (Crystal Chem Cat.#90080 and #90030, respectively). Estradiol (Cayman Chemical Cat.# 582251), testosterone (Cayman Chemical Cat.# 582701), FSH (Cayman Chemical Cat.#500710), LH (in house, as described ^113^) and AMH (Mybiosource Cat.# MBS268824) were measured in plasma samples obtained during the metestrus phase of the estrous cycle using relevant ELISA kits. In addition, LH pulsatility was assessed from the tail tip blood samples taken at 10-min intervals over 6 hours from 4-month-old females, as described previously ^114^.

### Histological analysis of ovaries and fat pads

Ovaries and gonadal fat pads collected from WT and *Gw* females (2, 4 and 6 months of age) were fixed in 4% formalin overnight, dehydrated with a series of increasing concentrations of ethanol solutions and cleared with xylene. The dehydrated tissues were embedded in paraffin wax and cut into 10μm sections. Then, the sections were rehydrated with decreasing concentrations of ethanol and stained with either haematoxylin and eosin (Sigma), Masson’s trichrome (Sigma), or oil red O stain (Sigma). For ovary sections, the follicles, corpora lutea, and cysts were counted in every 5^th^ sagittal section under a Leica DM2000 histology microscope, using previously described classification systems ^115, 116^: 1) Primordial follicle, consisting of one layer of flattened granulosa cells surrounding the oocyte; 2) Primary follicle, including one to two complete layers of cuboidal granulosa cells; 3) Secondary follicle, with an oocyte surrounded by more than one layer of cuboidal granulosa cells with no visible antrum; 4) Antral follicle, with an oocyte surrounded by multiple layers of cuboidal granulosa cells and containing one or more antral spaces, possibly with a cumulus oophorus and thecal layer; 5) Atretic follicle, with a follicle that enters a degenerative process without ovulation; 6) Corpora lutea, exclusively made of luteal cells that are relatively homogenous in terms of their size and distribution; and 7) Follicular cysts, with a thin wall composed of up to four layers of granulosa cells, without signs of luteinization, and filled with pale fluid or blood with or without cell debris/degenerating oocytes. Adipocyte size within gonadal fat pads was determined by measuring the surface area using ImageJ software.

### Hypothalamus micro-dissection and isolation/culture of hypothalamic cells

To isolate specific hypothalamic nuclei, freshly dissected brains from WT and *Gw* females (2 weeks or 4 months of age) were immediately cut into 1mm-thick sagittal sections in ice-embedded rodent brain matrix (ASI Instruments). The sections containing the arcuate nucleus and ventromedial/dorsomedial hypothalamus were identified and microdissected under a Leica M125 stereomicroscope according to key visible anatomical structures ^117^.

To evaluate the percentage of RFP-positive cells and expression levels of the *Gata4p[5kb]-RFP* transgene, freshly dissected hypothalamus from heterozygous and homozygous *Gw* mice (1 month of age) was minced into small pieces and subjected to enzymatic digestion in EMEM medium supplemented with 0.4mg/ml collagenase I, 1.3 mg/ml dispase II, and 0.1mg/ml DNase I. Each tissue piece was then gently triturated and incubated with gentle agitation at 37 °C for 30 min. RFP-positive and negative cells were recovered by fluorescence-activated cell sorting (FACS) using a BD FACSJazz cell sorter (BD Biosciences).

To establish primary cultures of hypothalamic neurons and astrocytes from 1-month-old heterozygous and homozygous *Gw* mice, we adapted previously described protocols for embryonic and early postnatal cortical neural cells ^118, 119^. Briefly, hypothalamus was freshly dissected in ice-cold Hank’s balanced salt solution (HBSS), minced into small pieces, and incubated for 15 minutes at 37°C in dissociation medium consisting of HBSS supplemented with 0.4mg/ml collagenase I, 1.3 mg/ml dispase II, and 0.1mg/ml DNase I. Samples were then gently triturated, starting with a 1000μl micropipette followed by a 200μl micropipette (25-30 times for each), until uniform cellular dissociation was achieved. The resulting cell suspension was filtered through a 70μm strainer, and cells were plated on poly-D-Lysine Gibco Cat.# A3890401) coated 8-chamber slides at a density of 50,000 cells/chamber. These cells were then pre-incubated at 37°C and 5% CO_2_ for 2h in Neurobasal medium (Gibco, Cat.# 10888-022) supplemented with 2% B27 (Gibco, Cat.# 17504-044), 1X Glutamax+ (Gibco, Cat.# 35050-061), 10% fetal bovine serum, 5% horse serum, and 1% penicillin/streptomycin, followed by culture for 7 days under the same conditions but without serum, until analysis via immunofluorescence (see below)

### Immunostaining

For immunofluorescence of fixed hypothalamic cells (on glass coverslips) and 10µm-thick paraffinized ovarian sections (on glass slides), cells/tissues were permeabilized for 2h in blocking solution (10% fetal bovine serum and 1% Triton X-100, in PBS), and then sequentially incubated with specific primary (overnight, at 4°C) and relevant secondary (at room temperature for 2h) antibodies diluted in same blocking solution (see Table S3 for detailed information about antibodies used). All sections were mounted with either DAPI or Hoechst as nuclear stain. A Nikon A1 confocal microscope equipped with NIS-Elements C software (Nikon) was used to acquire images, which were then analyzed using ImageJ. ER stress levels were quantified by counting the number of positive granulosa and theca cells, within growing follicles, and dividing them by the overall number of growing follicles in each ovary.

For immunostaining of hypothalamic nuclei, frozen brains were cut into 40µm coronal sections on a freezing sliding microtome (Leica SM2000 R) and mounted onto glass slides, which were then sequentially incubated in blocking solution (10% normal horse serum/0.3% Triton X-100/0.1% NaAzide) for 1h followed by specific primary (overnight, at 4°C) and relevant secondary (at room temperature for 1h) antibodies diluted in same blocking solution (see Table S3 for details). For immunohistochemistry, slides were finally stained with avidin-biotin horseradish peroxidase/3,3’-diaminobenzidine and counterstained with methyl green, using standard procedures. For immunofluorescence, slides were counterstained with DAPI. Images were taken using an Eclipse TE200 microscope (Nikon), an EMCCD ImageM digital camera (Hamamatsu), and iVision software (BioVision). Representative images were processed for viewing using ImageJ.

### Western blotting

Whole cell lysates of freshly dissected ovaries and hypothalamus from WT and *Gw* mice (4 months of age) were prepared in ice-cold RIPA buffer (150mM NaCl, 50mM Tris pH 8, 1mM EDTA, 1.0% Triton X-100, 0.1% SDS, 0.5% sodium deoxycholate) supplemented with cOmplete protease inhibitor cocktail (Roche). Resulting protein extracts (50µg) were then separated by SDS-PAGE, transferred onto polyvinylidene difluoride membrane, and processed for Western blotting. To this end, membranes were first blocked for 2 h at room temperature in TBST buffer (25 mM Tris·HCl pH 7.5, 150 mM NaCl, 1% Tween 20) supplemented with 5% non-fat dried milk (Carnation) and then incubated overnight at 4°C with primary antibodies diluted in TBST containing 1% BSA (see Table S3 for all details about antibodies used, including dilution factor). Membranes were then washed three times in TBST and incubated with the appropriate horseradish peroxidase-conjugated secondary antibody in TBST containing 5% non-fat dried milk. Immunoreactive bands were revealed by enhanced chemiluminescence (Immobilon western, Millipore Cat.# WBKLS0500) using the Fusion FX7 visualizing system (Montreal-BioTech). Quantification was done using ImageJ tools for gel analysis.

### RT-qPCR analysis

Total RNA from dissected hypothalamus nuclei was extracted using the RNeasy mini kit (Qiagen Cat.#74134), according to the manufacturer’s instructions. For each sample, 250-500 µg of total RNA was reverse transcribed using the SuperScript III kit (Invitrogen Cat.# 18080-044), according to the manufacturer’s protocol. Quantitative PCR was performed using a Bio-Rad C1000 thermal cycler (CFX96 Real Time System). Each amplification reaction mixture (20 μl) contained 10 ng cDNA equivalent to reverse transcribed RNA, 500 nM forward and reverse primers, 10 μl SensiFAST SYBR no-ROX Mix (Bioline Cat.# BIO-98005). Cycling conditions were: 1 cycle at 95 °C for 2 min, 40 cycles at 95 °C for 5 s followed by 60 °C for 30 s, and 1 final cycle at 65 °C for 5 s. SYBR green fluorescence was measured at the end of the annealing and extension phases of each cycle. Relative gene expression was calculated using the 2^-ΔΔCT^ method, using the house-keeping gene *Hprt1* for normalization. All primers used are shown in Table S2.

### Statistical analysis

Statistical analysis was performed using GraphPad Prism version 10.1.2. Two-tailed Welch’s *t*-tests were used for comparisons between two groups. When comparing between more than two groups, one-way or two-way ANOVA followed by either Dunnet’s or Tukey’s multiple-comparison tests were performed. Unless otherwise indicated, data are presented as scatter blot with mean ± SEM. *P*-values lower than 0.05 were considered significant.

## ACKNOWLEDGEMENTS

The authors thanks David W. Silversides (Université de Montréal) in whose lab the *Gata4* reporter mouse lines were generated, the Transgenic and Genome Editing core of the CERMO-FC research center for generation of the CRISPR knockout mouse lines and help with the induced ovulation studies, Ferdinand Roelfsema (Leiden University) for help with LH pulsatility analysis, and Elisabet Stener-Victorin (Karolinska Institutet) for thoughtful discussion about this project and critical reading of the manuscript. This study was not supported by any dedicated grants, being mainly funded by the UQAM Research Chair on Rare Genetic Diseases help by NP. SAN was also supported by postdoctoral fellowships from the Fonds de la Recherche du Québec – Santé (FRQS) and the Réseau Québécois en Reproduction (RQR), while ME was supported by a doctoral scholarship from the RQR.

## AUTHOR CONTRIBUTIONS

Conceived the study: NP, CM, RSV. Supervised the study: NP, CM. Designed the experiments: SAN, ME, NP, CM, RSV, DJB, SW. Performed the experiments: SAN, ME, OS, TC, GB, KFB, FG, FB, XZ, JS. Analyzed the data: SAN, ME, NP, CM, RSV, XZ, LS, DJB, JS, SW. Contributed reagents/materials/analysis tools: NP, CM, RSV, DJB, SW. Wrote the paper: SAN, ME, NP. Provided critical feedback: KFB, DJB, JS, SW, CM, RSV. All authors read and approved the final manuscript.

## DATA AVAILABILITY STATEMENT

All data generated or analyzed during this study are included in this article and its supplementary information files.

