## Supplementary material for "Brain-specific *Gata4* downregulation in *Greywick* female mice models the metabolic subtype of polycystic ovary syndrome": Fig.S

### Supplementary information

**Figure S1. Adult *Gw* males and *Gm10800/Gm10801* double-mutant females have normal fertility rates and weight.**

**Figure S2. A subset of *Gw* female mice kept on the C57BL/6N background is subfertile, but do not gain extra-weight with age.**

**Figure S3. Estrous cycle profile and cyst/follicle counts of *Gw* female mice at 2 and 4 months of age.**

**Figure S4. Levels of circulating FSH and AMH hormones are not significantly affected in adult *Gw* female mice.**

**Figure S5. Puberty is delayed in female *Gw* offspring.**

**Figure S6. Supplemental metabolic data for adult *Gw* females are consistent with the metabolic subtype of PCOS.**

**Figure S7. Copy number-dependent *Gata4p[5kb]-RFP* transgene expression in the hypothalamus of *Gw* female mice.**

**Figure S8. The total number of ER$\alpha$-positive cells in the Arc and GnRH neurons in the POA are not significantly affected in adult *Gw* female mice.**

**Table S1. Percentage of RFP-positive cells in the hypothalamus of *Gw* mice at 2, 4 and 8 weeks of age.**

**Table S2. List of all primers used in this study.**

**Table S3. List of primary antibodies used in this study.**

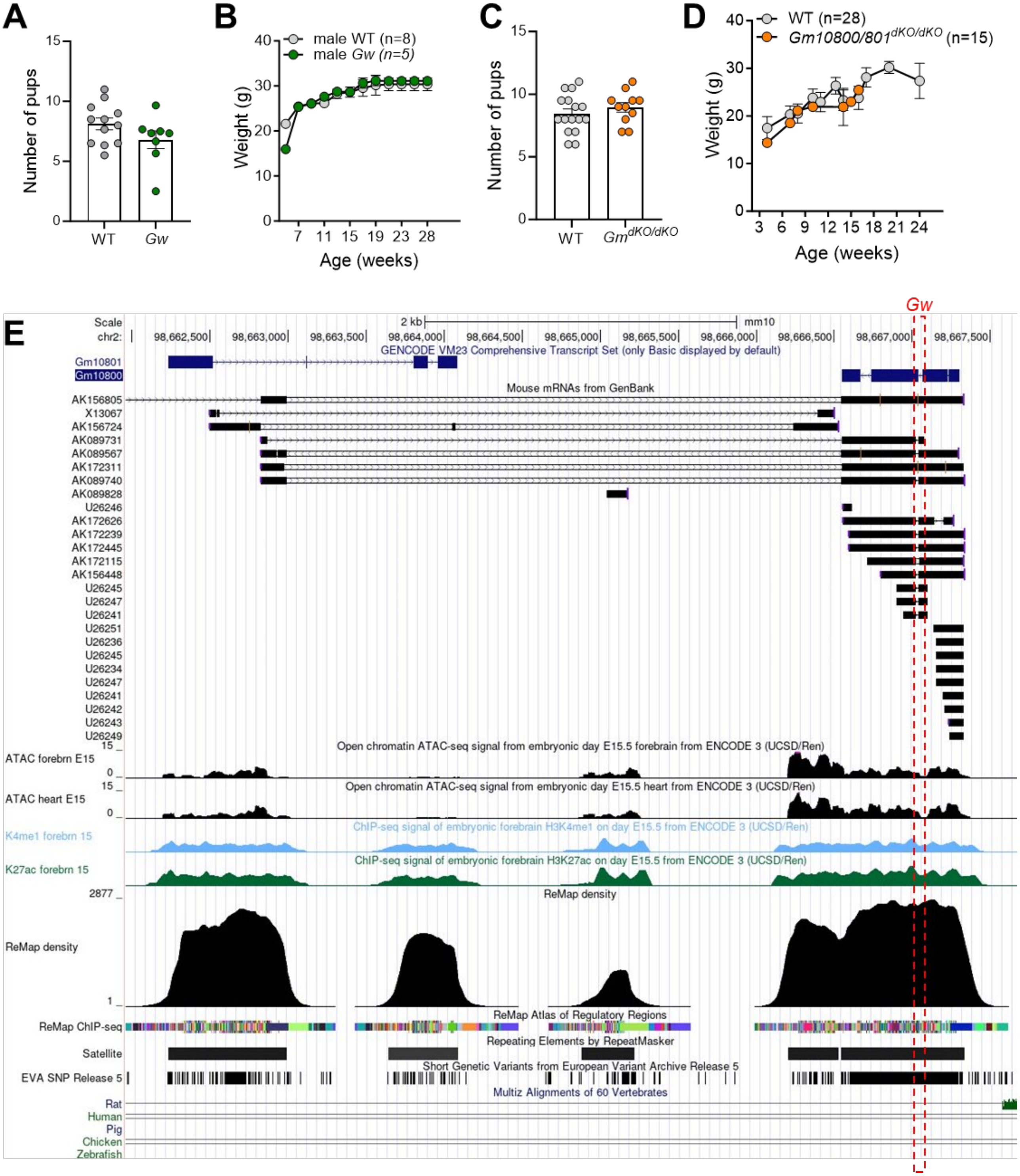


**Figure S1. Adult *Gw* males and *Gm10800/Gm10801* double-mutant females have normal fertility rates and weight. A)** Average number of pups per litter for WT females, after breeding with either WT or *Gw* males (n=8-12 females per group, 2-3 litters per female). **B)** Weight gain as a function of male age (*Gw vs* WT; n=5-8 males per group). **C)** Average number of pups per litter for *Gm10800/801*$^{dKO/dKO}$ and WT females (2- to 6-month-old), after breeding with WT males (n=11-16 females per group, 2-3 litters per female). **D)** Weight gain as a function of female age (*Gm10800/801*$^{dKO/dKO}$ *vs* WT; n=15-28 females per group). **E)** Screenshot from the UCSC genome browser showing key genomic features within and around both *Gm10800* and *Gm10801*. Same tracks than those displayed in Fig.2D are shown, with additional inclusion of the “Mouse mRNAs from Genbank” track showing multiple bidirectionally transcribed RNA species of variable lengths.

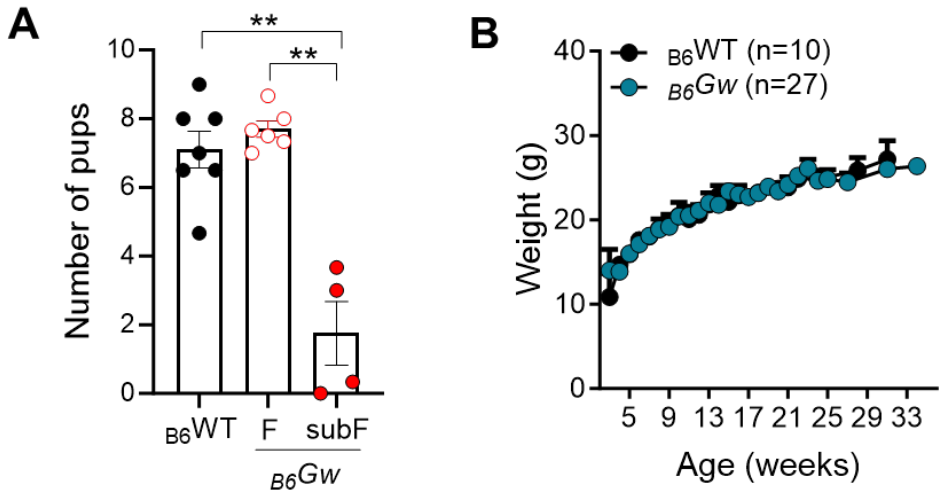


**Figure S2. A subset of *Gw* female mice kept on the C57BL/6N background is subfertile, but do not gain extra-weight with age. B)** Average number of pups per litter for adult *Gw* and WT females in C57BL/6N background (2- to 6-month-old), after breeding with WT males (n=4-7 females per group, 2-3 litters per female). F, fertile; subF, subfertile. **C)** Weight gain as a function of female age (*Gw vs* WT in C57BL/6N background; n=10-27 females per group). ** $p < 0.01$; two-tailed Welch's *t*-tests.

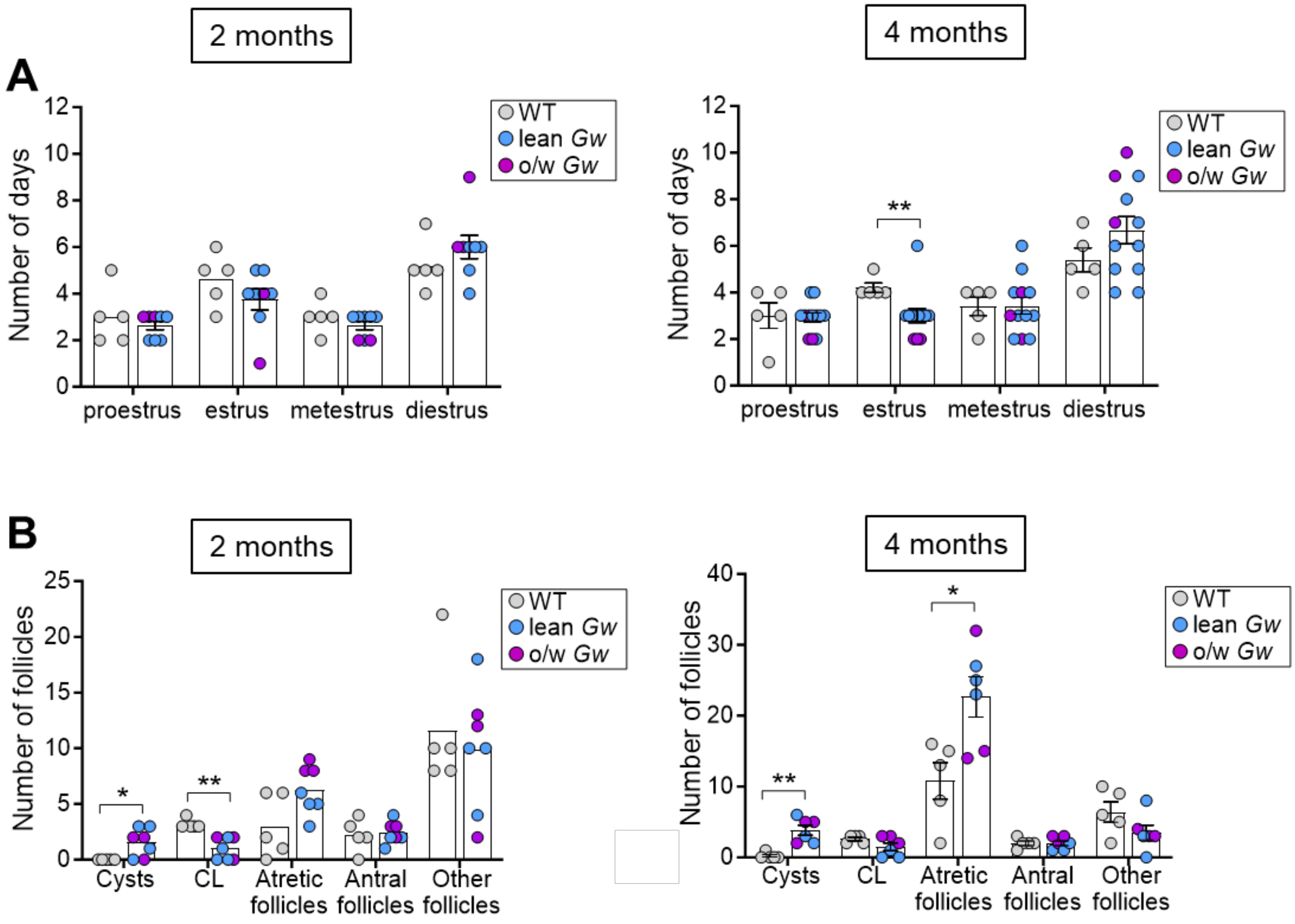


**Figure S3. Estrous cycle profile and cyst/follicle counts of *Gw* female mice at 2 and 4 months of age. A)** Number of days spent in each phase of the estrus cycle per period of 16 days in 2-month-old (left panel) and 4-month-old (right panel) mice (n=3-9 females per group). **B)** Average number of cysts, corpora lutea (CL), atretic follicles, antral follicles and other follicles counted on ovary cross-sections of 2-month-old (left panel) and 4-month-old (right panel) mice (n=3-5 females per group). * $p < 0.05$, ** $p < 0.01$; two-tailed Welch's *t*-tests.

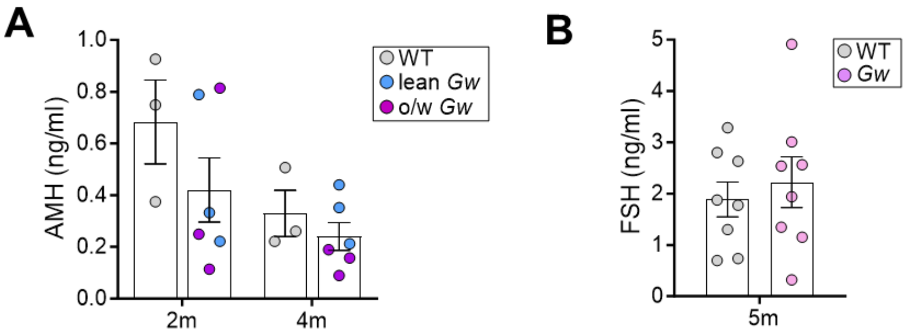


**Figure S4. Levels of circulating FSH and AMH hormones are not significantly affected in adult *Gw* female mice. A)** Circulating plasma levels of AMH in 2- and 4-month-old WT and both lean and overweight (o/w) *Gw* female mice at metestrus (n=3 mice per group). **B)** Circulating plasma levels of FSH in 5-month-old WT and *Gw* female mice at metestrus (n=8 mice per group).

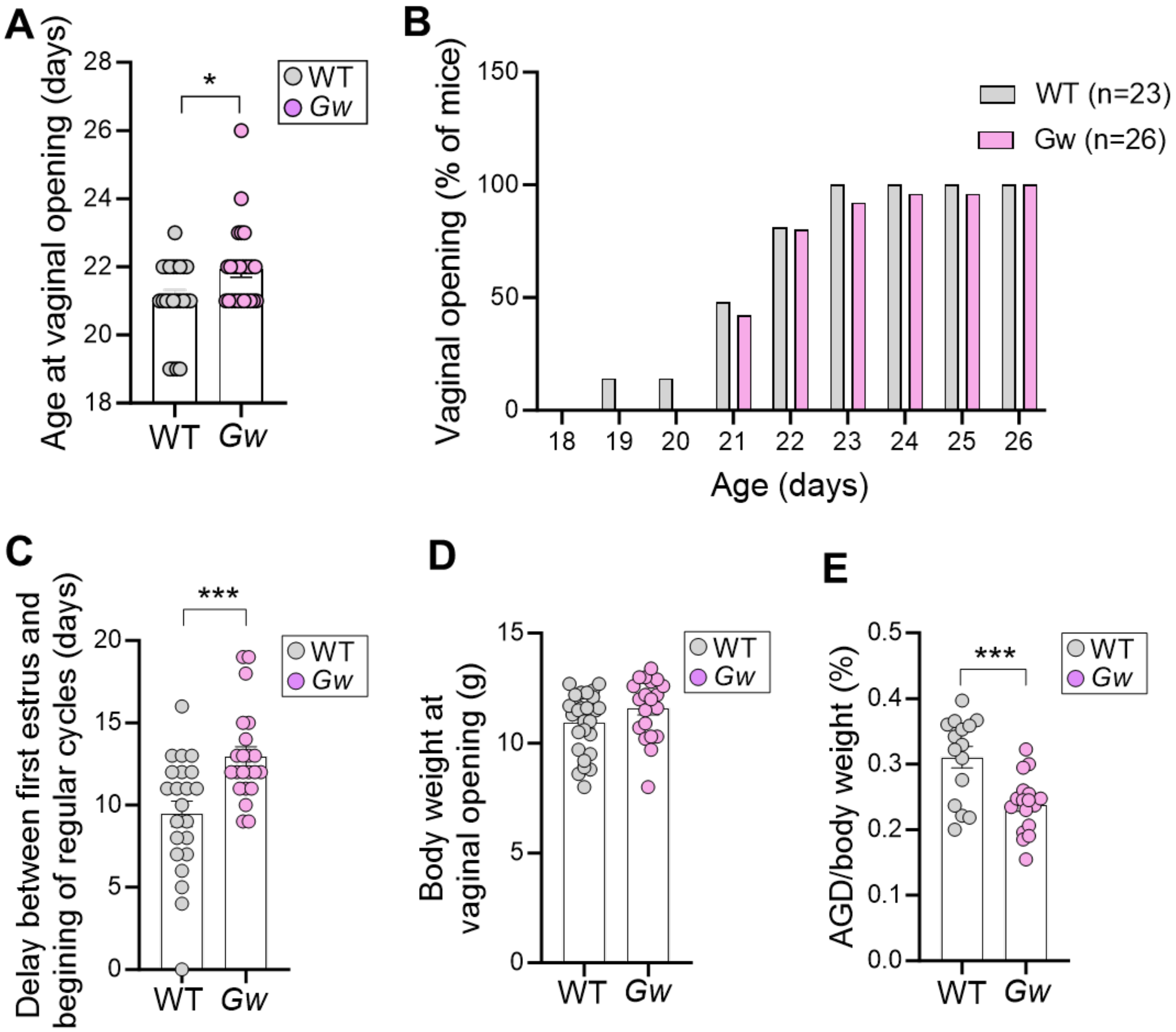


**Figure S5. Puberty is delayed in female *Gw* offspring. A)** Age of WT and *Gw* female mice at vaginal opening (n=23-26 mice per group). **B)** Percentage of female mice with opened vagina from postnatal day 18 to postnatal day 30 (n=23-26 mice per group). **C)** Delay between first estrus and the beginning of regular estrous cycles in WT and *Gw* female mice (n=22 mice per group). **D)** Body weight of WT and *Gw* female mice at vaginal opening (n=20-27 mice per group). **E)** Anogenital distance (AGD) of WT and *Gw* female mice at postnatal day 20 (n=15-17 mice per group). * $p < 0.05$, *** $p < 0.001$; two-tailed Welch's *t*-tests.

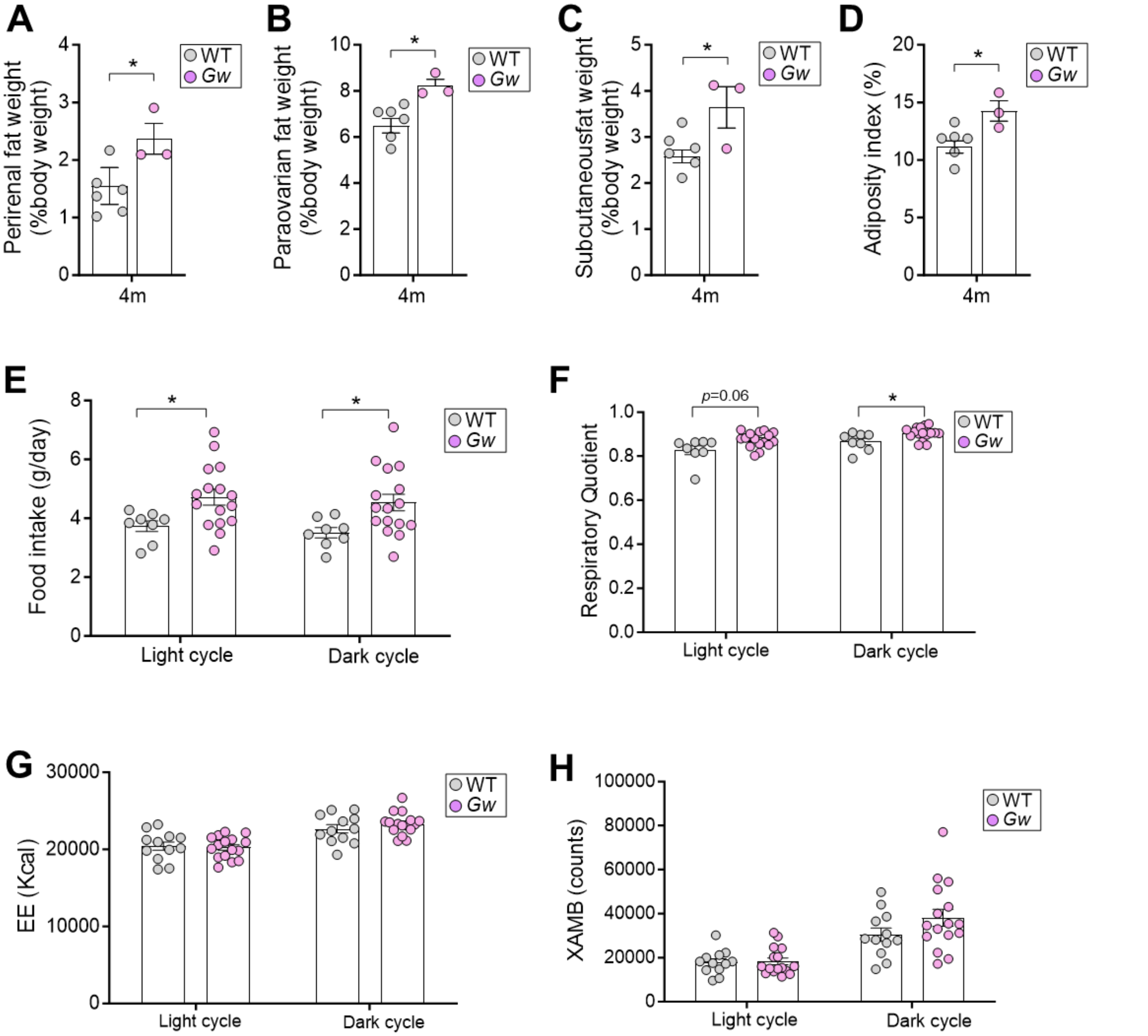


**Figure S6. Supplemental metabolic data for adult *Gw* females are consistent with the metabolic subtype of PCOS. A-C)** Weight of perirenal (A), paraovarian (B), and subcutaneous (C) fat pockets expressed as percentage of body weight in 4-month-old WT and *Gw* female mice (n=3-6 mice per group). **D)** Adiposity index expressed as percentage of body weight in 4-month-old WT and *Gw* female mice (n=3-6 mice per group). **E-H)** Light and dark cycle CLAMS analysis of average food intake (E), respiratory quotient (F), energy expenditure (EE) (G), and X-ambulatory movement (XAMB) (H) over a 48h time interval in 4-month-old WT and *Gw* female mice (n=8-16 mice per group). * $p < 0.05$; two-tailed Welch's *t*-tests.

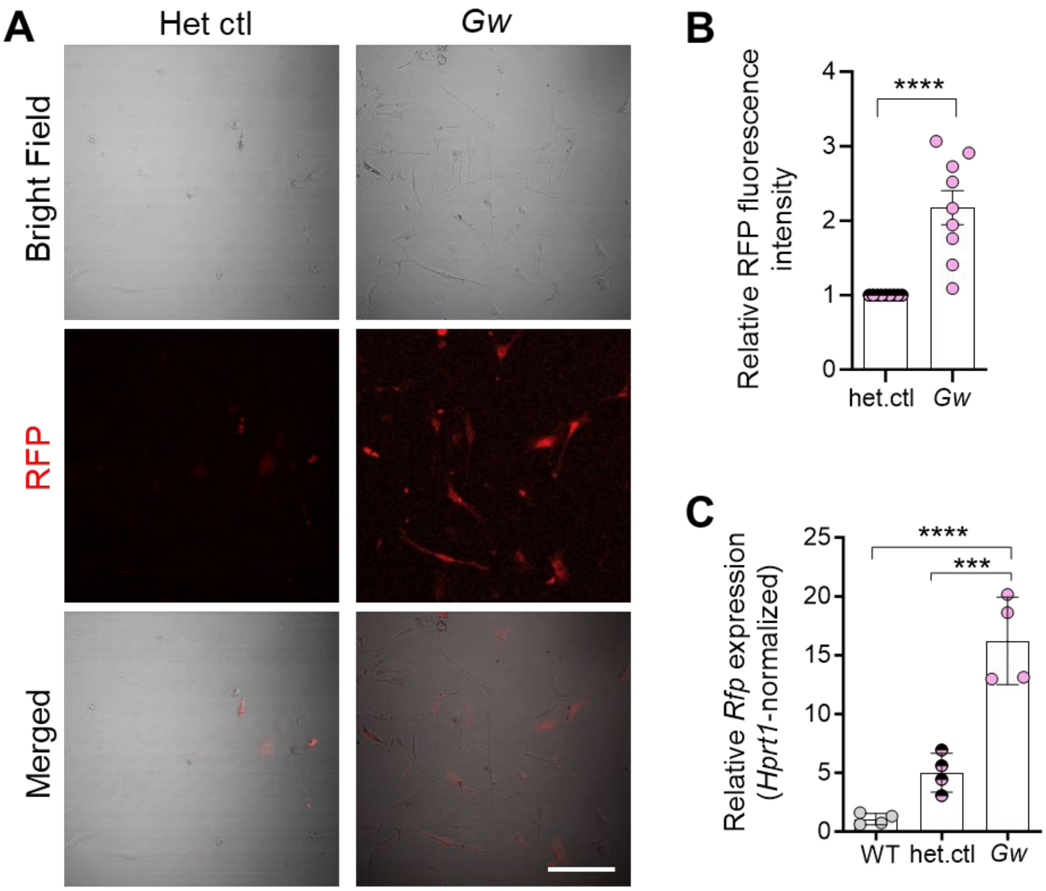


**Figure S7. Copy number-dependent *Gata4p[5kb]-RFP* transgene expression in the hypothalamus of *Gw* female mice. A)** Representative confocal images of RFP fluorescence in unfixed hypothalamic neural cells cultured for 7 days after microdissection/dissociation from 1-month-old heterozygous (het. ctl) and homozygous *Gw* female mice (n=3 mice per group). **B)** Quantification of RFP fluorescence intensity using images such as those displayed in panel A (n=3 mice per group, 3 fields of view per sample). **C)** RT-qPCR analysis of *Rfp* expression levels in the hypothalamus of 1-month-old WT and both heterozygous (het. ctl) and homozygous *Gw* female mice (n=4 mice per group). *** $p < 0.001$, **** $p < 0.0001$; two-tailed Welch's $t$-tests.

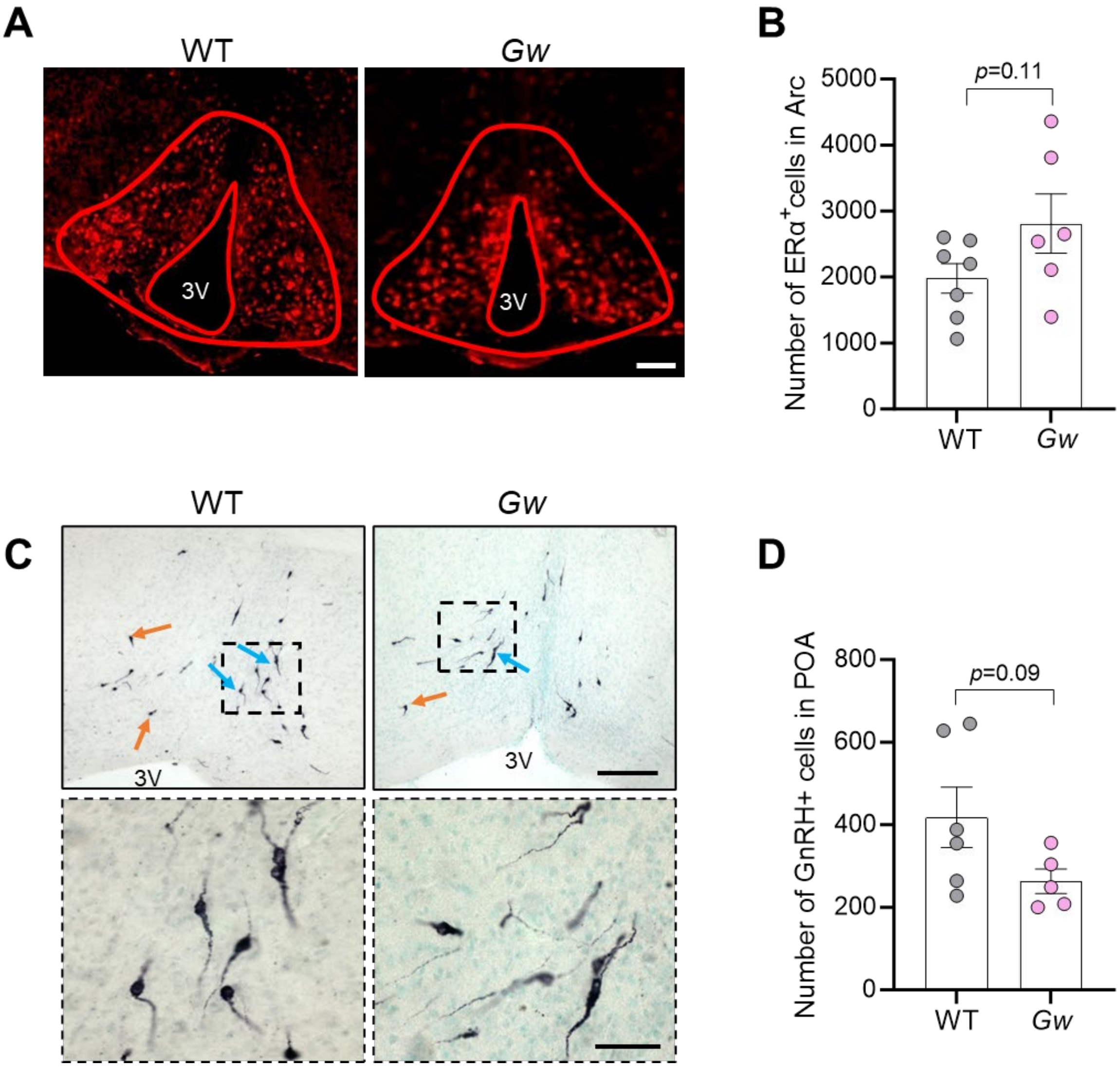


**Figure S8. The total number of ERα-positive cells in the Arc and GnRH neurons in the POA are not significantly affected in adult *Gw* female mice. A)** Representative photomicrographs of coronal brain sections from 4-month-old WT and *Gw* mice showing ERα+ cells in the Arc at low magnification (10x). 3V, third ventricle. Scale bar, 100 μm. **B)** Quantitative analysis of the total number of ERα+ cells per Arc of WT and *Gw* mice, using images such as those displayed in panel A (n=6-7 mice per group). **C)** Representative photomicrographs of coronal brain sections from 4-month-old WT and *Gw* mice showing GnRH+ neurons in the POA at low (10x, upper panels) and high (40x, lower panels) magnification. Orange and blue arrows point to unipolar and bipolar cells, respectively. 3V, third ventricle. Scale bar, 100 μm (upper panels) and 25 μm (lower panels). **D)** Quantitative analysis of the total number of GnRH neurons per POA of WT and *Gw* mice, using images such as those displayed in panel C (n=5-6 mice per group). Two-tailed Welch's *t*-test.

**Table S1. Percentage of RFP-positive cells in the hypothalamus of *Gw* mice at 2, 4 and 8 weeks of age.**

| Age | Het. ctl *Gw* female mice (n=3 mice per time point) | *Gw* female mice (n=3 mice per time point) |
|---|---|---|
| *2 weeks* | 17 277 cells (0.44%) | 12 579 cells (0.81%) |
| | 12 265 cells (0.51%) | 11 499 cells (0.59%) |
| | 14 268 cells (0.63%) | 19 393 cells (0.97%) |
| *4 weeks* | 13 409 cells 0.29%) | 14 743 cells (0.35%) |
| | 12 522 cells (0.35%) | 20 495 cells (0.60%) |
| | 10 240 cells (0.29%) | 15 800 cells (0.45%) |
| *8 weeks* | 2 977 cells (0.04%) | 10 899 cells (0.14%) |
| | 3 903 cells (0.03%) | 10 812 cells (0.086%) |
| | 4 818 cells (0.03%) | 10 645 cells (0.098%) |

**Table S2. List of all primers used in this study.**

| | |
|---|---|
| ***Greywick* genome-transgene boundaries** | |
| **Ch2_Fwd** | 5'-GCACACTGAAGGACCTGGAATATG |
| **RFP_Rev** | 5'-CCTAATATTCTAATAGCAGCCATAAAATACTCC |
| **Tyrosinase_Fwd** | 5'-GCTGTTTGGCTTTTTTTCAGACCC |
| **Ch2_ Rev** | 5'-CATATTCCAGGTCCTTCAGTGTGC |
| ***Greywick* genotyping** | |
| **Gw allele_Fwd** | 5'-GGGCTGTCATCTCACTATGGGCA |
| **_Rev** | 5'-TGATTATGTCCCCATGACTGTCAG |
| ***Gm10800* KO genotyping** | |
| **Gm10800_Fwd** | 5'-ATTCATTCATAAGTGGATAGTGGC |
| **_Rev1** | 5'-TAACTCCTGGGCTCAAGCAGTC |
| **_Rev2** | 5'-TGACGAAATCATCTGTTTCCAAAG |
| **RT-qPCR** | |
| **Hprt1_Fwd** | 5'-TCAGTCAACGGGGGACATAAA |
| **_Rev** | 5'-GGGGCTGTACTGCTTAACCAG |
| **Rfp_Fwd** | 5'-TGT CCC CCC AGT TCC AGT A |
| **_Rev** | 5'-GTT GTG GGA GGT GAT GTC CA |
| **Gata4_Fwd** | 5'-CTCTGGAGGCGAGATGGGAC |
| **_Rev** | 5'-CGCATTGCAAGAGGCCTGGG |
| **Esr1_Fwd** | 5'-AGGACCACATCCACCGTGTC |
| **_Rev** | 5'-GGGATTCTCAGAACCTTTCGG |
| **Pgr_Fwd** | 5'-CATACCTTAACTACCTGAGG |
| **_Rev** | 5'-GATTAAGCAGATCTTCTGAGG |
| **Gnrh1_Fwd** | 5'-GCTCCAGCCAGCACTGGTCCTA |
| **_Rev** | 5'-TGATCCACCTCCTTGCCCATCTCTT |
| **Kiss1_Fwd** | 5'-GGAGAAGGACCTGTCGACCT |
| **_Rev** | 5'-GACGGCAGCATTGCTTTTAT |
| **Nknb_Fwd** | 5'-GCCATGCTGTTTGCGGCTG |
| **_Rev** | 5'-CCTTGCTCAGCACTTTCAGC |
| **Pdyn_Fwd** | 5'-CCCTCTAATGTTATGGCGGA |
| **_Rev** | 5'-AGAGACCGTCAGGGTGAGAA |
| **Pomc_Fwd** | 5'-ATGCCGAGATTCTGCTACAGT |
| **_Rev** | 5'-TCCAGCGAGAGGTCGAGTTT |
| **Agrp_Fwd** | 5'-CGGAGGTGCTAGATCCACAGA |
| **_Rev** | 5'-AGGACTCGTGCAGCCTTACAC |
| **Npy_Fwd** | 5'-CTCCGCTCTGCGACACTACA |
| **_Rev** | 5'-AATCAGTGTCTCAGGGCTGGA |
| **Lepr (long)_Fwd** | 5'-GGTCTCAGAGCACCCAGGTA |
| **_Rev** | 5'-TGGATAAACCCTTGCTCTTCATC |
| **Lepr (short)_Fwd** | 5'-CAATGTGGATCAGGATCAACC |
| **_Rev** | 5'-AGCCAGAACTGTAACAGTGTG |

**Table S3. List of primary antibodies used in this study.**

| Antibody name (host and clonality) | Source | Catalog number (RRID) | Dilution factor (method) |
|---|---|---|---|
| *ERα (C1355)* *(rabbit polyclonal)* | Upstate (Millipore) | 06-935 (AB_310305) | 1:2500 (IF) |
| *GAPDH* *(mouse monoclonal)* | Santa Cruz | SC-32233 (AB_627679) | 1:5000 (WB) |
| *GATA4* *(mouse monoclonal)* | Santa Cruz | SC-25310 (AB_627667) | 1:1000 (WB)<br>1:200 (IF) |
| *GnRH* *(rabbit polyclonal)* | Home-made (Wray lab) | SW-1 (AB_2629221) | 1:15000 (IHC) |
| *IRE1α – phospho S724* *(rabbit polyclonal)* | Abcam | ab48187 (AB_873899) | 1:500 (IF) |
| *NeuN* *(rabbit polyclonal)* | Abcam | ab104225 (AB_10711153) | 1:500 (IF) |
| *RFP* *(chicken polyclonal)* | Novus Biologicals | NBP197371 (AB_11139267) | 1:500 (IF) |
| *RFP* *(mouse monoclonal)* | Thermo Fisher Scientific | MA5-15257 (AB_10999796) | 1:1000 (WB) |
| *S100β* *(rabbit monoclonal)* | Abcam | ab52642 (AB_882426) | 1:500 (IF) |
